# *miR-125-chinmo* pathway regulates dietary restriction dependent enhancement of lifespan in *Drosophila*

**DOI:** 10.1101/2020.09.01.277772

**Authors:** Manish Pandey, Sakshi Bansal, Sudipta Bar, Nicholas S. Sokol, Jason M. Tennessen, Pankaj Kapahi, Geetanjali Chawla

## Abstract

Dietary restriction (DR) extends healthy lifespan in diverse species. Age and nutrient-related changes in the abundance of microRNAs (miRNAs) and their processing factors have been linked to organismal longevity. However, the mechanisms by which they modulate lifespan and the tissue-specific role of miRNA mediated networks in DR dependent enhancement of lifespan remains largely unexplored. We show that two neuronally enriched and highly conserved microRNAs, *miR-125* and *let-7* mediate the DR response in *Drosophila melanogaster*. Functional characterization of *miR-125* demonstrates its role in neurons while its target *chinmo* acts both in neurons and the fat body to modulate fat metabolism and longevity. Proteomic analysis revealed that Chinmo exerts its DR effects by regulating expression of *FATP, CG2017, CG9577, CG17554, CG5009, CG8778, CG9527* and *FASN1*. Our findings identify *miR-125* as a conserved effector of DR pathway and open up the avenue for this small RNA molecule and its downstream effectors to be considered as potential drug candidates for treatment of late-onset diseases and biomarkers for healthy aging in humans.

## INTRODUCTION

Aging is characterized by a progressive decline in physiological function, which leads to an increased risk of chronic degenerative diseases and disabilities(Harman, 2003). Deregulated nutrient signaling is one of the key hallmarks of aging, and restricting nutrient intake or dietary restriction (DR) has been shown to enhance health and longevity in most species(Fontana & Partridge, 2015; Kapahi, Kaeberlein, & Hansen, 2017; Klass, 1977; Lin et al., 2002; Lopez-Otin, Blasco, Partridge, Serrano, & Kroemer, 2013; McCay, Crowell, & Maynard, 1989). More significantly, DR delays age-related pathologies such as diabetes, cardiovascular diseases, cancer and neurodegenerative disorders(Anson et al., 2003; Klebanov, 2007; Kraus et al., 2019; Maswood et al., 2004; Mattison et al., 2012; Mattson & Wan, 2005; Wang et al., 2005). Emerging evidence from diverse model systems has implicated micro-ribonucleic acids (miRNAs) as critical components of signaling pathways that modulate lifespan by regulating mRNA turnover and translation(Boehm & Slack, 2005; Chawla, Deosthale, Childress, Wu, & Sokol, 2016; Liu et al., 2012; Verma, Augustine, Ammar, Tashiro, & Cohen, 2015). These 19-24 nucleotides long, single-stranded RNAs function by directing effector complexes to target mRNAs(Bartel, 2018). This recruitment of the miRNA ribonucleoprotein complexes (miRNPs) is facilitated by interactions between the miRNA and its target and results in silencing of the target mRNA(Guo, Ingolia, Weissman, & Bartel, 2010; Kim, 2005). Since, the interaction of a miRNA and its target occurs by imperfect base-pairing interactions, a single miRNA can target several mRNAs in a given context. Thus, these evolutionary conserved and dosage sensitive effectors possess the key attributes to facilitate the complex metabolic reprogramming that occurs during dietary restriction. While studies in the *C. elegans*, mammalian cell culture, mouse and primate model systems have reported regulation of miRNAs and their targets upon DR, there is no evidence to indicate whether the DR mediated expression changes of the miRNAs and their downstream targets occur in the same tissue and whether modulating miRNA and their target mRNAs can modulate lifespan extension by DR(Mercken et al., 2013; Pandit, Jain, Kumar, & Mukhopadhyay, 2014; Schneider et al., 2017; Zhang et al., 2019).

Here we report that nutrient restriction in *D. melanogaster* upregulates *let-7-Complex* miRNAs (*miR-100, let-7* and *miR-125*). Furthermore, *let-7* and *miR-125* loss of function mutations dampen the dietary restriction dependent lifespan extension. The DR phenotype associated with loss of *miR-125* is due to the derepression of its target, *<u>Ch</u>ronologically <u>In</u>appropriate <u>Mo</u>rphogenesis (chinmo*). Our analysis reveals that *chinmo* codes for a nutrient regulated transcription factor, and its upregulation in the nervous system results in altered fat metabolism. Consistent with the *miR-125* loss of function DR phenotype, increasing the dosage of human *miR-125* in the fat body increased longevity. In summary, we have identified a conserved miRNA that mediates the effects of DR by promoting tissue-tissue communication demonstrating its potential as a DR mimetic agent.

## RESULTS

### Dietary restriction dependent upregulation of *let-7* and *miR-125* increases lifespan

To examine whether *let-7-Complex* miRNAs mediate the effects of dietary restriction (DR), we investigated whether DR affects the expression of these miRNAs in wild type (*w_1118_*) and *let-7-C* hypomorphic (*let-7-C_hyp_*) mutants. Wild type and mutant flies were exposed to dietary restriction or *ad libitum* (AL) conditions for 7, 20 or 30 days, and quantitative reverse transcription polymerase chain reaction (qRT-PCR) analyses was performed with RNA extracted from whole animals. Under AL conditions, young *let-7-C_hyp_* mutants displayed lower levels of *miR-100* (36 ± 16.5%), *let-7* (29 ± 6.9%) and *miR-125* (70 ± 18%) relative to the wild type strain in AL conditions (Figure 1B). Increasing the dietary restriction for 20 days led to an increase in the levels of *miR-100* (*W_1118_*: 223 ± 73%; *let-7-C_hyp_*: 140 ± 16.7%), *let-7* (*w_1118_*: 170 ± 19.7%; *let-7-C_hyp_*: 190 ± 20%) and *miR-125* (*w_1118_*: 250 ± 49%; *let-7-C_hyp_*: 172 ± 40%) in both wild type and *let-7-C_hyp_* mutants. Though, increasing DR for 30 days led to an increase in the levels of all three miRNAs in both the strains, the *let-7-C_hyp_* mutant expressed significantly lower levels of *miR-100* (AL: 46 ± 4.3%; DR: 95 ±7.1%), *let-7* (AL: 56.2 ± 5.25%; DR: 129 ± 15.1%) and *miR-125* (AL: 56 ± 3.8%; DR: 75 ± 3.4%) relative to *w_1118_* flies on AL diet. To examine whether the increase in *let-7-C* miRNAs was required for lifespan extension upon DR, we examined the survival of *w_1118_* and *let-7-C_hyp_* mutants fed an AL and DR diet (Figure 1C, D). Wild type (*w_1118_*) flies fed a DR diet (blue line) lived significantly longer than wild type flies that were fed an AL diet (red line). DR increased median lifespan by 50% in *w_1118_* flies and by 0% in the *let-7-C_hyp_* mutants (Figure 1D and Supplementary Table 1A). These data indicated that one or more of the *let-7-C* miRNAs were required for DR mediated lifespan extension.

To determine the contribution of *miR-100, let-7* and *miR-125* in DR mediated enhancement of lifespan, we measured survival of *miR-100, let-7* and *miR-125* mutant flies reared on AL or DR diet (Figure 1E-H). *let-7-C_null_* Rescue (See Supplementary Table 7 for genotype) and Δ*miR-100* female flies fed a DR diet (blue line) lived significantly (*w_1118_* DR: 47% increase median lifespan; Δ*miR-100* DR: 18.75% increase in median lifespan) longer than flies that were fed an AL diet (Figure 1E, F and Supplementary Table 1B). Δ*let-7* mutants exhibited a significantly dampened lifespan extension when fed a DR diet. Though, a 33% increase in median lifespan was observed, the DR fed flies had a 4.7% decrease in maximum lifespan compared to the AL fed flies (Figure 1G and Supplementary Table 1B). In contrast, Δ*miR-125* mutants failed to exhibit lifespan extension when fed a DR diet and a 0% increase in median lifespan was observed for Δ*miR-125* flies that were fed a DR diet (Figure 1H and Supplementary table 1B). These data confirmed the requirement of *miR-125* and *let-7* in DR dependent extension of lifespan.

**Figure 1.**
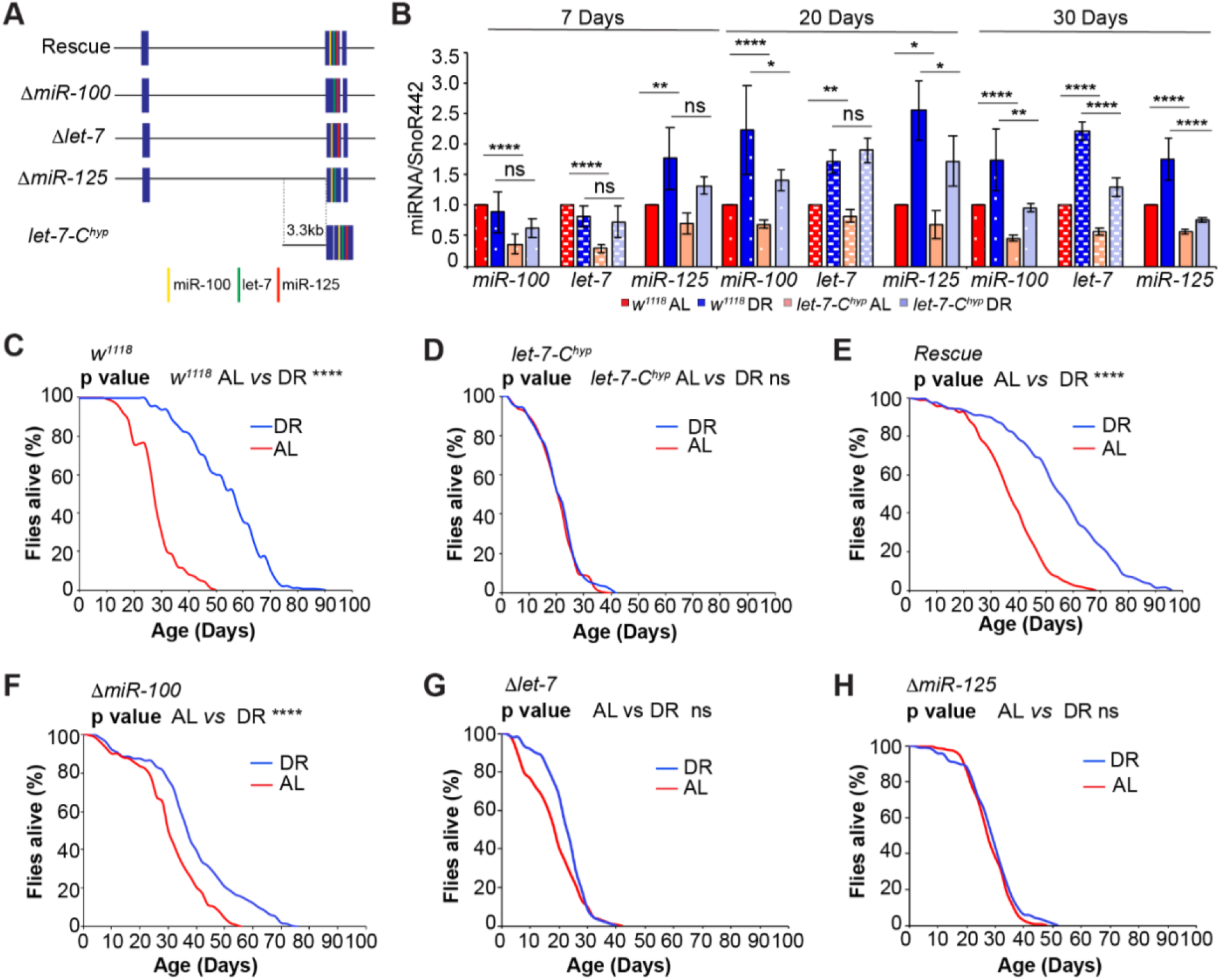
*let-7* and *miR-125* are required for dietary restriction (DR) dependent enhancement of lifespan. (A) Schematic of the *let-7-Complex* (*let-7-C*) transgenes. The 18kb Rescue *let-7-C* construct encodes all three miRNAs; *miR-100, let-7* and *miR-125*. The *ΔmiR-100, Δlet-7* and *ΔmiR-125* lack miR-100, *let-7* and *miR-125*, respectively. The *let-7-C* hypomorph encodes the *let-7-Complex* cDNA driven by a 3.3kb promoter fragment in a *let-7-C* null mutant. The *let-7-C* rescue line encodes a single copy of the 18kb *let-7-C* locus in a *let-7-C* null mutant. (B) Quantitative RT-PCR of *miR-100* (bars with white dots), *let-7* (bars with white dashes) and *miR-125* (solid filled bars) in wild type (*w_1118_*) (red and dark blue) and *let-7-C_hyp_* (pink and light blue) flies that were fed *Ad libitum* (AL) (red and pink) or DR diet (dark blue and light blue) for 7 days, 20 days or 30 days. Expression levels were normalized to SnoR442. Values are mean ± SD, n≥ 3. (C) *w_1118_* flies show a significant increase in lifespan upon dietary restriction (DR, blue line) as compared to *w_1118_* flies that were fed an *“ad libitum”* (AL) diet (red line). p value < 10-10. (D) *let-7-C_hyp_* mutants do not display a DR-dependent increase in lifespan (compare red and blue curves). p value= 0.2219. (EH) Compared to the rescue flies (E) and *ΔmiR-100* (F) flies, *Δlet-7* and *ΔmiR-125* flies displayed a significantly dampened DR-dependent lifespan extension upon DR (blue curve) when compared to flies that were fed an AL diet (red curve). For statistical analysis for comparison of survival curves, p values were calculated with log rank test. p value * <0.05.

### *miR-125* regulates dietary restriction by repressing *chinmo*

To test whether the DR dependent phenotypes of the *miR-125* loss of function mutant were due to lack of post-transcriptional silencing of its previously validated target *chinmo*, we measured survival of strains in which dosage of *chinmo* was reduced genetically by a *let-7-C* Gal4 driven *chinmo_RNAi_* transgene or a *chinmo_1_* loss of function mutant in the *let-7-C* rescue flies and in *miR-125* mutants (Figure 2 A-B and Supplementary Figure 1A-B) (Chawla et al., 2016; Wu, Chawla, & Sokol, 2020; Wu, Chen, Mercer, & Sokol, 2012; Zhu et al., 2006). Lowering *chinmo* levels suppressed the DR-dependent lifespan phenotypes of *miR-125* mutants (Compare Figure 1H with Figure 2A–2B and Supplementary Table 2A-2B). Though, reducing *chinmo* specifically in *let-7-C* expressing (*let-7-C Gal4*) cells was able to increase the median lifespan by 25% (compared to 0% in *miR-125* mutants), a 75% increase in median lifespan was observed when *chinmo* levels were reduced genetically (*chinmo_1_*) in all *chinmo* expressing cells (Figure 2B and Supplementary Table 2A-2B). These data demonstrate a role for *chinmo* in dietary restriction dependent lifespan extension by *miR-125*. Furthermore, these data indicated that *chinmo* played a wider role in regulating DR-dependent lifespan extension which is not just limited to *miR-125* cells.

**Figure 2.**
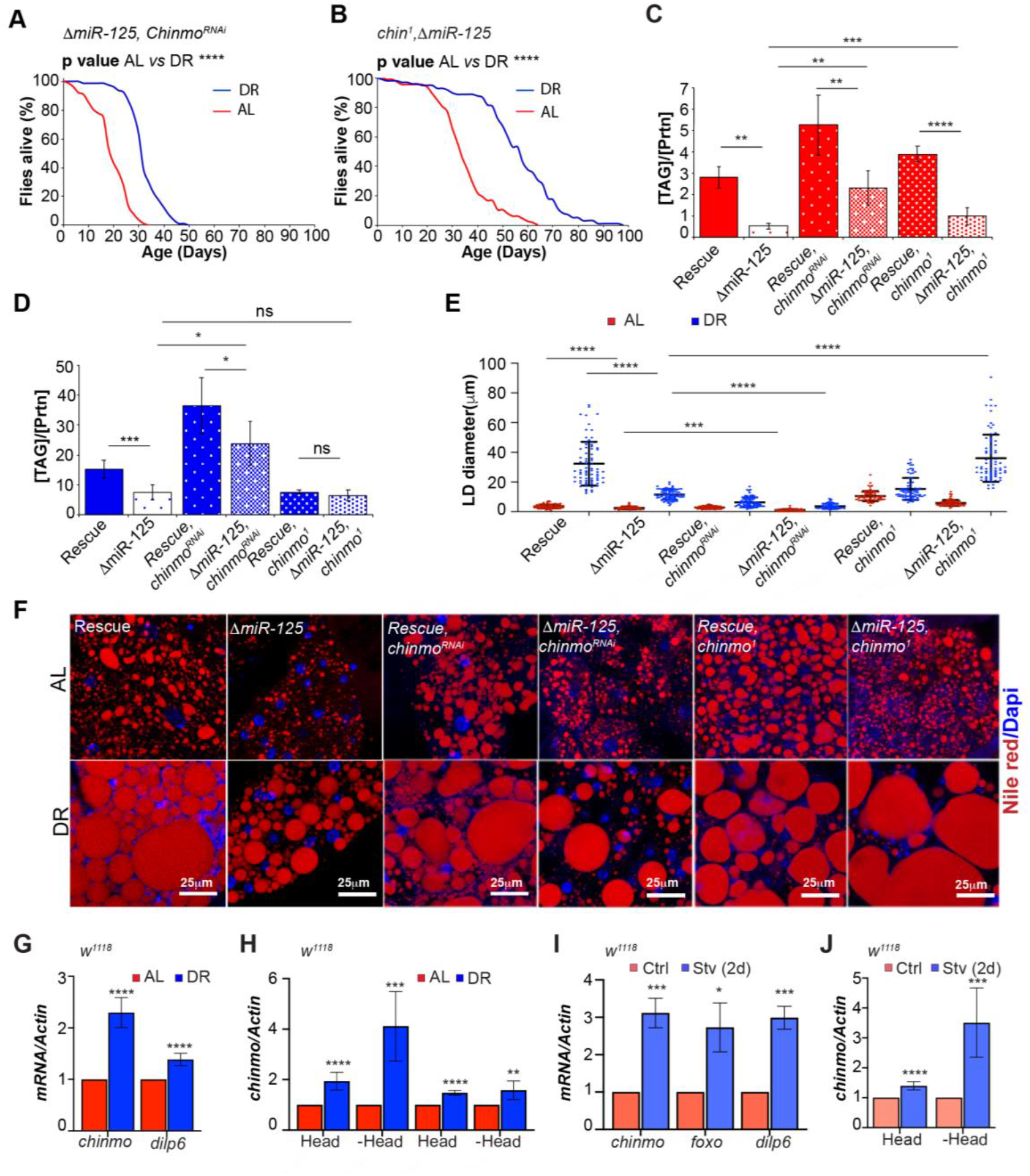
Reducing *chinmo (ΔmiR-12, chinmo_RNAi_ and ΔmiR-125, chinmo_1_)* levels in *miR-125* mutants suppresses the loss of DR mediated increase in lifespan extension and decrease in triglyceride levels. (A) Δ*miR-125, UAS chinmo_RNAi_* flies show a 25% increase in median lifespan upon dietary restriction (DR, blue line) as compared to *ΔmiR-125, UAS chinmo_RNAi_* flies that were fed an *“Ad libitum”* (AL) diet (red line). p value < 10^-8^. (B) *ΔmiR-125, chinmo_1_* flies display a 75% increase in DR-dependent increase in median lifespan (compare red and blue curves). p value<10-10. For statistical analysis for comparison of survival curves, p values were calculated with log rank test. p value * <0.05. (C, D) Quantitation of triglyceride (TAG) stored levels in Rescue, *ΔmiR-125*, Rescue, *UAS chinmo_RNAi_, ΔmiR-125, UAS chinmo_RNAi_. Rescue, chinmo_1_* and *ΔmiR-125, chinmo_1_*, AL (Red bars) and DR (Blue bars) fed 20 d old flies. The bars represent mean ± SD, n ≥ 5, p value was calculated with two tailed t-test. Significance levels: ***p<0.001. (E, F) Fat bodies/abdomens of female flies were dissected and stained for the content and diameter of lipid droplet (LD) (red are lipid droplets stained with Nile red and blue is Dapi). Scale bar, 25μm. (E) Quantitation of lipid droplet (LD) diameter in (F). Quantitation of 15 largest LDs in 5 samples per condition. Error bars represent ± SD and **** p <0.0001. (G, H) DR induces expression of *chinmo* in wild type (*w_1118_*) flies. (G) RT-PCR quantitation of fold change in *chinmo* and *dilp6* mRNA levels in *w_1118_* flies that were an AL (Red bars) or DR (Blue bars) for 30 days. (H) The DR-dependent increase in *chinmo* is higher in peripheral tissues as compared to the head tissue. RT-PCR quantitation of *chinmo* mRNA in head and decapitated body tissue of *w_1118_* flies that were fed an AL or DR diet for 10 days or 20 days. (I, J) Starvation induces expression of *chinmo* in wild type flies. (J) A significantly higher increase in *chinmo* mRNA is seen in tissues other than head upon exposure of *w_1118_* flies to starvation for 2 days. Expression levels were normalized to *Actin5c*. Values are mean ± SD, n≥ 3

DR imposed by restricting yeast in the diet enhances lipid content (Bradley & Simmons, 1997; Katewa et al., 2012) and this increase in lipid content enhances lipid turnover under DR and is required for the DR dependent lifespan extension in *Drosophila* (Katewa et al., 2012). Since, *miR-125* mutants displayed a DR phenotype, we quantitated the levels of total triglycerides (TAG) in whole bodies of rescue and *miR-125* mutants that were exposed to AL or DR diet for 20 days. In contrast to the rescue strain, the *miR-125* mutants displayed a significant drop in TAG levels in both AL (81 ± 4.6%) and DR (50 ± 16.4%) conditions (Figure 2C-D). In a parallel experiment, we also examined lipid droplets in the fat body of adult flies by staining with Nile red. Consistent with the TAG analysis, the *miR-125* mutants displayed a significant drop in the diameter of lipid droplets compared to the rescue strain in both AL (35% decrease in the median) and DR (59% decrease in median) diet (Figure 2E-F). Thus, our analysis demonstrates that *miR-125* regulates the shift in lipid metabolism upon DR. Consistent with the lifespan data, reducing *chinmo* dosage in *ΔmiR-125* flies resulted in an increase in the TAG levels and lipid droplet diameter (Figure 2E–2F). However, a much greater increase in TAG levels was observed in *let-7-C Gal4> ΔmiR-125, UAS chinmo_RNAi_* (4.4 ± 1.55-fold increase in AL and 3.18 ± 0.98-fold increase in DR relative to *ΔmiR-125*) flies as compared to the *ΔmiR-125, chinmo_1_* flies (1.89 ± 0.76-fold increase in AL and 0.87 ± 0.24-fold increase in DR) (Figure 2 C, D). These differences could arise due to other unknown *miR-125* targets operating in the DR pathway or due to *mir-125* independent regulation of *chinmo* upon DR in cells that do not express *miR-125*. We then examined whether nutrient-dependent regulation of *chinmo* was linked with the DR dependent differences in the lifespan and fat metabolism. Wild type (*Canton S* or *w_1118_*) flies that were fed a DR diet for 30 days expressed ~2-fold higher levels of *chinmo* RNA than the AL fed flies (Figure 2G). This increase in expression was higher in tissues other than head (Figure 2H). The expression of *chinmo* mRNA also increased (~3-fold) upon starvation in *w_1118_* flies and a much greater increase was detected in peripheral tissues other than head (Figure 2I-J). Thus, implying that in addition to its regulation by *miR-125, chinmo* likely plays a DR dependent functional role in tissues other than the adult brain. Taken together, these data revealed that *chinmo* acts downstream of miR-125 to regulate lifespan extension upon DR.

### Overexpression of *chinmo* diminishes the DR-mediated lifespan extension

To gain an understanding of how tissue specific regulation of *chinmo* results in DR dependent lifespan extension, we measured survival of fruit flies that ectopically expressed *chinmo* in different tissues in adulthood under AL and DR conditions. Consistent with the *miR-125* single mutant phenotype, overexpression of *chinmo* using a ubiquitous *Daughterless* GAL4 gene switch (*Da GS*), fat body specific gene switch driver (*BL8151*), an intestinal specific (*5966*) or neuronal gene switch driver (*3X Elav GS*) resulted in a decrease in the DR dependent lifespan extension (Figure 3A, B, C, D) (*DaGS > UAS chinmo*: a 62.5% increase in median lifespan -RU and a 0% increase in +RU; *5966 > UAS chinmo*: a 41.66% increase in median lifespan in -RU and a 0% increase in median survival in +RU; *FBGS > UAS chinmo*: an 82.3% increase in DR in -RU and a 11% increase in median survival + RU; *3X ElavGS > UAS chinmo*: a 154% increase in median survival in -RU and a 14.2% increase in median survival in +RU). A significantly dampened DR mediated lifespan extension was observed when *chinmo* was overexpressed in the adult intestine (Figure 3B and Supplementary Table 3B). In addition, ectopic neuronal expression of *chinmo (3XelavGS*) also resulted in decreased TAG levels and a concomitant reduction in lipid droplet diameter in both AL and DR diets (Figure 3F-H). Consistent with the *miR-125* single mutant metabolic phenotypes (Figure 2 C-F), ectopic expression of *chinmo* in the adult fat body with an inducible fat body specific GAL4 driver (Supplementary Figure 2A-D) resulted in reduced TAG levels (Supplementary Figure 2B) and a decrease in lipid droplet diameter (Supplementary Figure 2C-D). Together, these data demonstrate a role for *miR-125* in DR pathways via post-transcriptional repression of *chinmo*.

**Figure 3.**
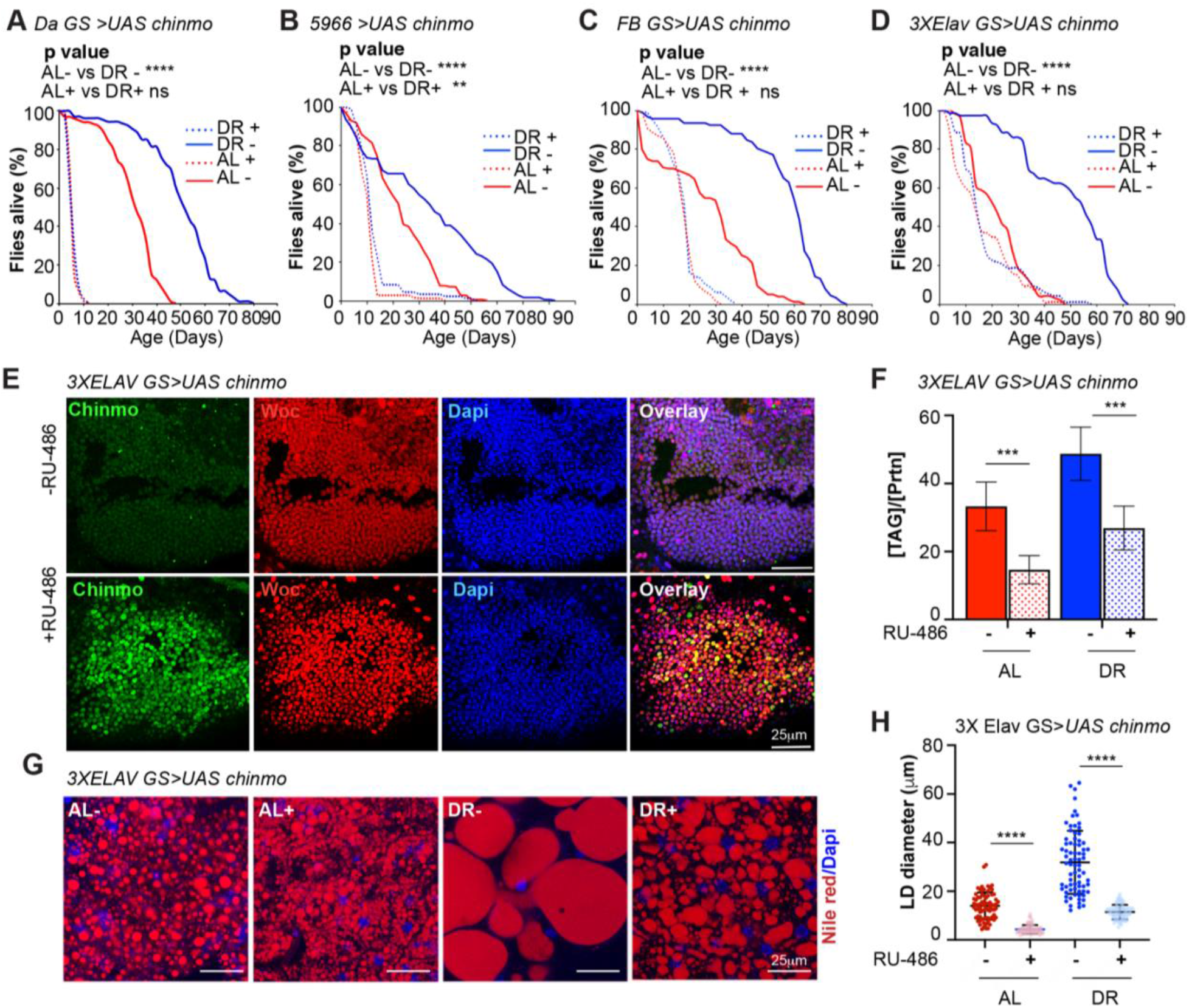
Ectopic expression of *chinmo* dampens the DR dependent lifespan increase and reduces fat metabolism. *UAS chinmo* was expressed in different adult tissues using the steroid (RU-486) inducible gene switch Gal4 drivers. (A) *DaGS > UAS chinmo* flies that were not fed RU-486 show a significant increase in lifespan upon dietary restriction (DR, blue line) as compared to *DaGS > UAS chinmo* flies that were fed an “*ad libitum”* (AL) diet (red line). RU-486 fed *DaGS > UAS chinmo* flies did not display a DR-dependent increase in lifespan (blue and red dotted lines). (B, C, D) *5966 > UAS chinmo, FB GS > UAS chinmo* and *3XElavGS > UAS chinmo* flies that were fed RU-486 displayed a significantly diminished lifespan extension upon DR (blue and red dotted lines) compared to the flies that were fed the solvent (blue and red solid lines). (E) Female flies that were fed an RU-486 supplemented diet for 5 days displayed increased levels of Chinmo in neuronal cells, as detected by Chinmo (green), Woc (red) and Dapi (blue) staining of dissected adult fly brains. (F) Quantitation of triglyceride (TAG) stored levels in AL-RU-486 (Solid red bars), AL + RU-486 (Red dotted bars), DR-RU-486 (Solid blue bars) and DR +RU-486 (Dotted blue bars) fed 20 d old *3XElavGS >UAS chinmo* flies. The bars represent mean ± SD, n ≥ 5, p value was calculated with two tailed t-test. Significance levels: ***p<0.001. (G) Fat bodies/abdomens of female flies were dissected and stained for the content and diameter of lipid droplet (LD) (red are lipid droplets stained with Nile red and blue is Dapi). Scale bar, 25μm. (H) Quantitation of lipid droplet (LD) diameter in (G). Quantitation of 15 largest LDs in 5 samples per condition. Error bars represent ± SD and **** p <0.0001.

### Non-autonomous role of Chinmo in dietary restriction

*Let-Complex* miRNAs are predominantly expressed in the adult nervous system however, our analysis has revealed that derepression of *chinmo* in the nervous system of *miR-125* mutants results in systemic changes in fat metabolism (Figure 3D-H) (Supplementary Figure 3). In order to determine the mechanistic basis for systemic regulation of fat metabolism by the *miR-125-chinmo* regulatory axis operating in the adult brain, we examined expression of Chinmo after inducing its expression with different gene switch Gal4 drivers (Figure 4A-H’). Immunohistochemistry was performed on dissected brain, ovary, gut and fat body of *DaGS > UAS chinmo, 5966 > UAS chinmo, FB GS > UAS chinmo* and *3X Elav GS > UAS chinmo* flies that were fed an AL or DR diets in presence and absence of RU-486. As expected, the Chinmo protein was detectable in all four tissues when the ubiquitous *Da GS* Gal 4 was used to drive its expression (Figure 4B, B’). Interestingly, Chinmo protein was also detectable in all four tissues when either the intestinal specific (*5966*) or neuronal (*3X Elav GS*) gene switch Gal4 were used to drive its expression (Figure 4D, D’-H, H’). However, Chinmo expression in the ovaries was restricted to the stalk when *5966 > UAS chinmo* flies that were fed RU-486 containing AL or DR diet. *FB GS > UAS chinmo* flies that were fed RU-486 containing food displayed expression of Chinmo in the fat body, gut and some in the stalks of ovaries. However, no Chinmo protein was detected in the brain (Figure 4F, F’). Quantitative RT-PCR analysis of *chinmo* mRNA in the dissected tissues of *3X Elav GS > UAS chinmo*, 5966> *UAS chinmo* and *FB GS > UAS chinmo* also indicated the presence of significantly higher levels of *chinmo* mRNA in tissues other than adult brain, gut and fat body, respectively. Thus, indicating the likelihood that increased levels of *chinmo* RNA was capable of diffusing to other tissues (Supplementary Figure 4).

**Figure 4.**
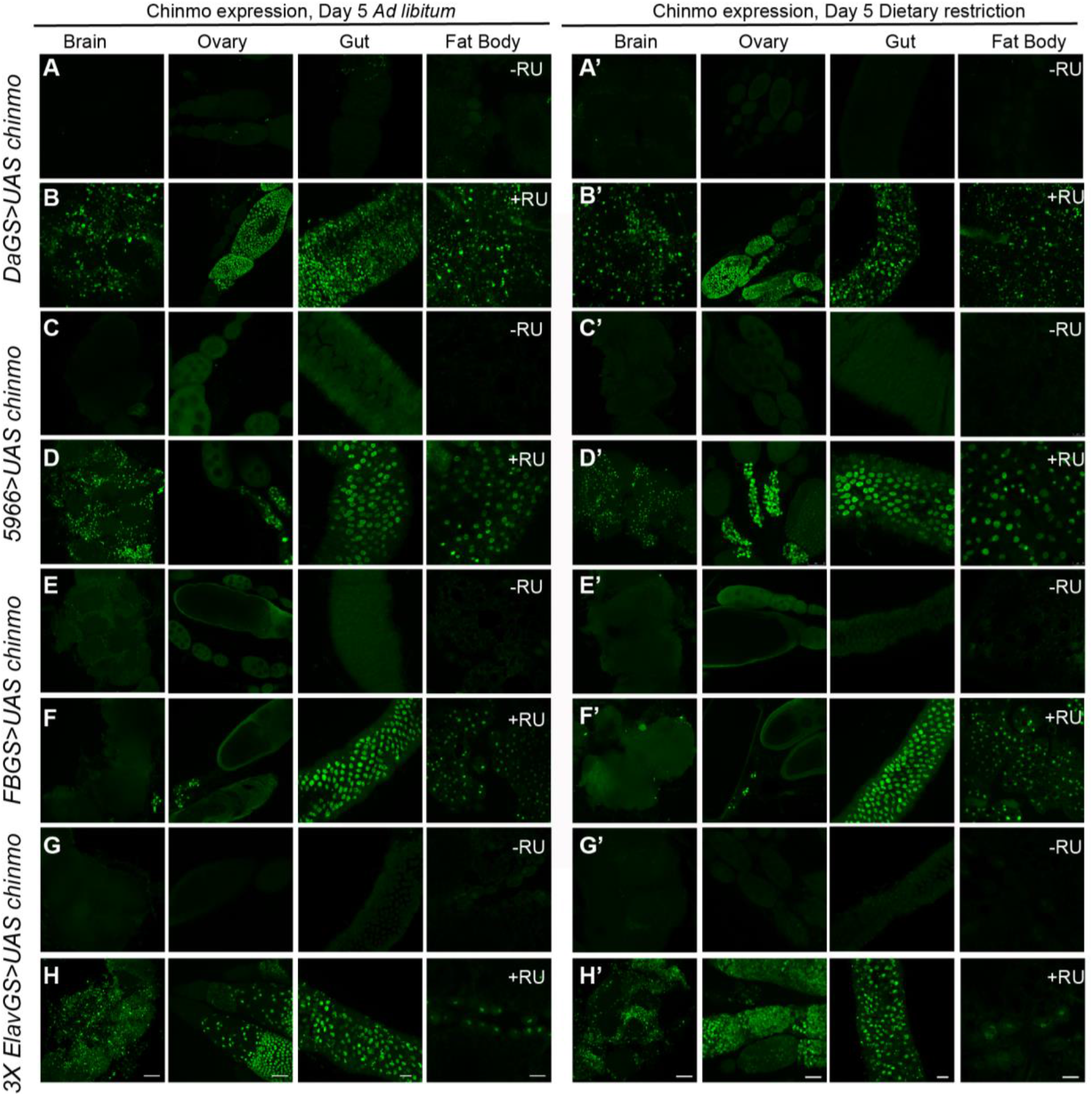
Ectopic expression of *chinmo* in adult neurons is not restricted to the brain tissue. *UAS chinmo* expression was driven in adult tissues in a drug inducible manner with a ubiquitously expressing *Da-GS* (4A, 4A’-4B, 4B’), *5966* (gut specific-GS) (4C, 4C’-4D, 4D’), fat body specific (*FB-GS*) (4E, 4E’-4F, 4F’) or neuronal (*3X Elav-GS*) gene switch GAL4 drivers (4G,4G’-4H, 4H’). (A-H’) Confocal images of dissected *Drosophila* tissues (adult brain, ovaries, gut and fat body) from fruit flies that were fed an AL diet (4A-H) or DR diet (4A’-H’) under uninduced (4A, 4C, 4E, 4G, 4A’,4C’, 4E’, 4G’) and steroid-induced (F, H, J, L, F’, H’, J’, L’) conditions. Scale bar, 50μm (brain, ovaries and fat body). Scale bar, 25μm (gut).

To confirm that the expression of Chinmo in peripheral tissue was not due to leaky expression of the Gal4 gene-switch drivers we drove expression of GFP using the *Da-GS, 5966, FB GS* and *3X Elav GS* drivers. Immunohistochemistry with GFP antibody confirmed the specificity of all four gene switch GAL4 drivers. As expected, GFP protein was detected in adult brain, ovaries, gut and fat body of *DaGS > UAS GFP* flies that were fed an AL+ RU or DR+ RU diet (Supplementary Figure 5B-B’). GFP was detected in the gut and fat body of *5966 > UAS GFP* and *FB GS > UAS GFP* flies upon RU treatment, although the fat body expression in *5966> UAS GFP* was much lower than that in the *FB GS> UAS GFP* indicating these two drivers have overlapping expression pattern (Supplementary Figure 5D, D’-F, F’). *3X ElavGS > UAS GFP* showed a slightly differential expression pattern on AL and DR diets (Supplementary Figure 5H, H’). The majority of GFP signal was observed in the brain and neurons innervating the fat body/abdominal wall. However, there were some cells in the gut that had GFP expression and this signal increased upon DR. Unlike *3XElav GS > UAS chinmo* flies, there was no GFP detected in ovaries of *3X ElavGS > UAS GFP* flies that were fed an AL diet but some GFP was detected in the stalks in flies that were fed a DR diet. These data indicated that Chinmo protein and/or mRNA was capable of diffusing to the other tissues and this non-autonomous expression of Chinmo mediated its effects on fat metabolism and lifespan.

### Knockdown of *chinmo* in the adult fat body enhances lifespan

To address the contribution of *chinmo* in DR dependent increase in lifespan, we examined the survival of flies expressing a *chinmo_RNAi_* transgene in the adult fat tissue (Figure 5A-B). A fat body (FB) gene switch Gal4 driver was used to overexpress *UAS-chinmo_RNAi_* in an inducible manner (Figure 5A). Knockdown of *chinmo* resulted in a 19% increase in median lifespan in flies that were fed an AL diet (Figure 5B). Since, the increase in lifespan upon DR was much lower than that of *FB GS* > *UAS-chinmo_RNAi_* flies that were fed an AL diet, it is likely that the lifespan extension mediated by reducing *chinmo* operates predominantly through the DR pathway. Reducing *chinmo* in the adult FB also resulted in an increase in stored triglyceride levels and an increase in lipid droplet diameter in flies that were fed an AL diet (Figure 5C-E). These data confirmed that *chinmo* functioned as an effector of DR mediated extension of lifespan.

**Figure 5.**
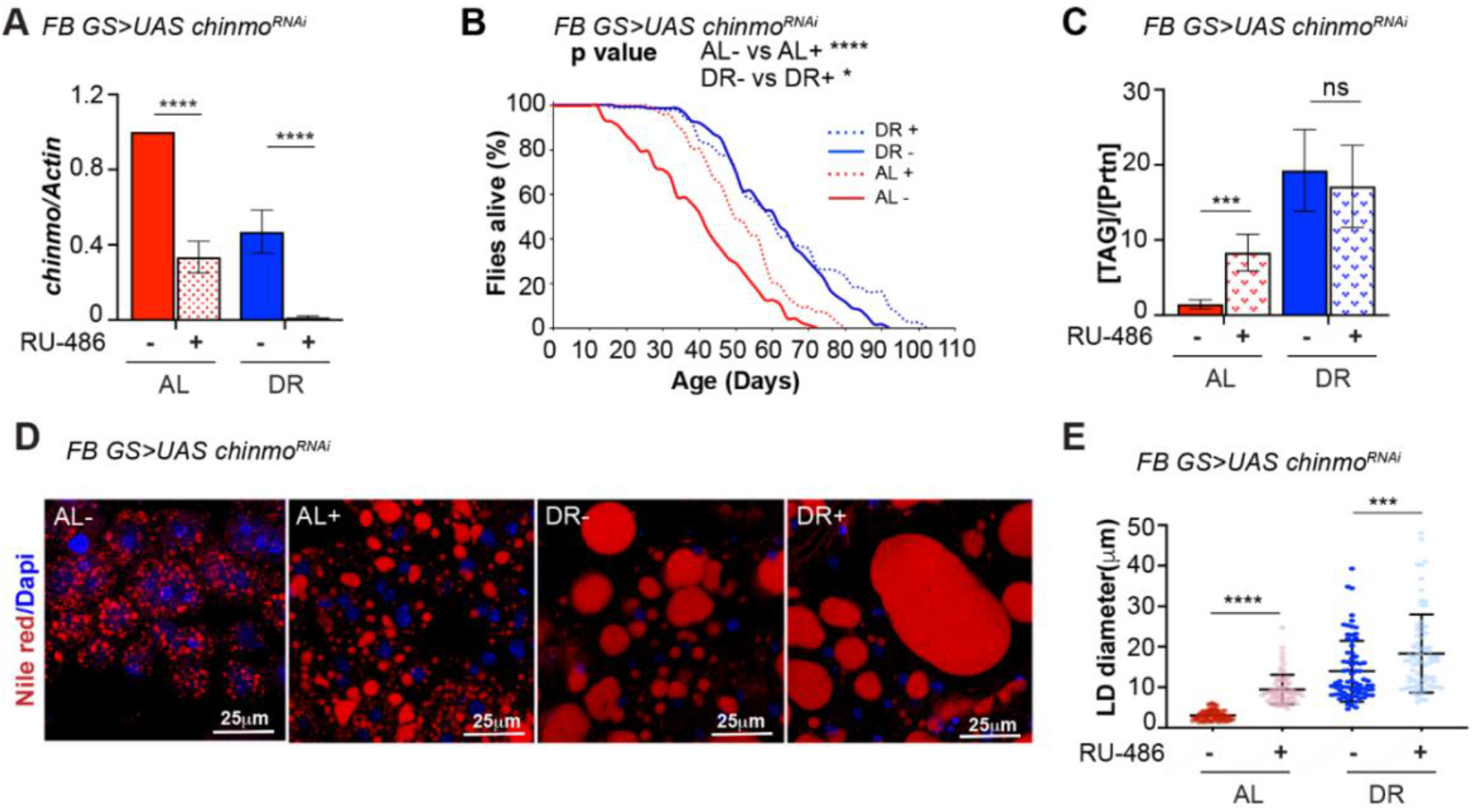
Reducing *chinmo* levels in the adult fat body increases lifespan and enhances lipid metabolism. (A) A transgene expressing a short hairpin to knockdown *chinmo* was expressed in the adult fat body using the steroid (RU-486) inducible gene switch Gal4 driver. (B) Knockdown of *chinmo* in the adult fat tissue resulted in a significant increase in the survival of flies that were fed an AL diet (compare red solid line with the red dotted line) (See Supplementary Table 3 for p values and median and maximum lifespan). (C) Quantitation of triglyceride (TAG) stored levels in AL-RU-486 (Solid red bars), AL + RU-486 (Red pattern bars), DR-RU-486 (Solid blue bars) and DR +RU-486 (Blue pattern bars) fed 20 d old *FB GS >UAS chinmo_RNAi_* flies. The bars represent mean ± SD, n ≥ 5, p value was calculated with two tailed t-test. Significance levels: ***p<0.001. (D) Fat bodies of *FB GS >UAS chinmo_RNAi_* female flies were dissected and stained for the content and diameter of lipid droplet (LD) (red are lipid droplets stained with Nile red and blue is Dapi). Scale bar, 25μm. (E) Quantitation of lipid droplet (LD) diameter in (D). Quantitation of 15 largest LDs in 5 samples per condition. Error bars represent ± SD and *** p <0.001.

### Chinmo downregulates expression of genes involved in fat metabolism

In order to identify potential targets of Chinmo that maybe responsible for the DR-dependent phenotypes of *miR-125* mutant, we performed semi quantitative proteomic analysis of extracts prepared from adult flies (whole animals) overexpressing *chinmo* specifically in adult neurons using the 3X *Elav GS* Gal4 driver (Figure 6A-E). Since the role of Chinmo as a repressor of gene expression is well established, we examined the downregulated biological processes to identify relevant direct downstream targets of Chinmo (Figure 6F). Proteins that were identified to be significantly downregulated were predominantly genes that were involved in metabolism (Figure 6F). We successfully validated the expression of seven fat metabolism genes (*FATP, CG2017, CG9577, CG17554, CG5009, CG8778, CG9527* and *FASN1*) that were identified through this proteomic analysis (Figure 6G-H) using RNA extracted from head tissue (Figure 6G) or decapitated fly tissue (Figure 6H) of *3XElavGS > UAS chinmo* flies. Consistent with the proteomics data, the mRNAs of all the fat metabolism genes were significantly downregulated in the head tissue (51-78% relative to control) and in the decapitated fly tissue (27-50% relative to control) of flies that ectopically expressed *chinmo* in adult neurons (Figure 6G-H). In order to test whether downregulation of the candidate fat metabolism genes was responsible for modulating lifespan, we measured survival of flies that expressed transgenes to knockdown *FASN1* and *FATP* specifically in the adult fat body (Figure 6I, J). Knockdown of *FASN1* resulted in a 14.7% decrease in median lifespan of flies that were fed an AL diet and a 31.8% decrease in median lifespan on DR diet (Figure 6I). Knockdown of FATP resulted in a 13% decrease in median lifespan upon DR (Figure 6J). Consistent with the RNAi data, overexpressing *UAS-Flag-FATP* specifically in the adult fat tissue increased median life span by 9.5% under DR conditions (Figure 6K). Taken together, these results indicated that ectopic expression of *chinmo* in the adult neurons causes decreased expression of fat metabolism genes in the fat tissue that resulted in a decreased lifespan.

**Figure 6.**
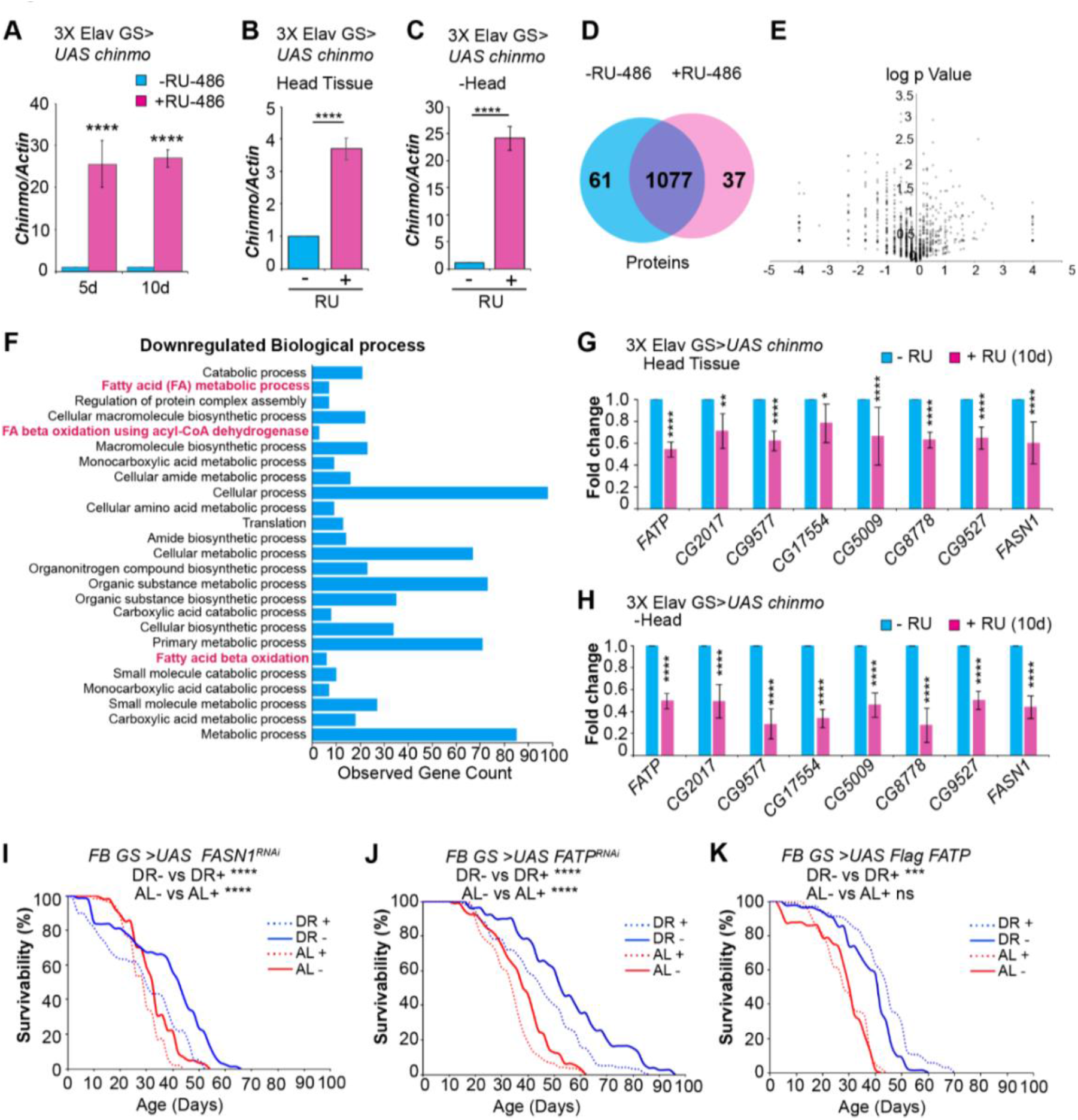
Ectopic expression of *chinmo* in adult neurons represses fat metabolism. (A) Quantitation of *chinmo* mRNA levels in RNA extracted from *3X ElavGS > UAS chinmo* flies that were fed a solvent (light blue bars) or RU-486 (pink bars) diet for 5 and 10 days. (B, C) Quantitation of *chinmo* mRNA levels in RNA extracted from head tissue (B) and decapitated body tissue (C) of *3X ElavGS > UAS chinmo* flies that were fed a solvent (blue bar) or RU-486 (pink bar) diet for 10 days. Expression levels were normalized to *Actin5c*. Values are mean ± SD, n≥ 3. (D) Venn diagram of 1175 proteins identified across the two groups (-RU-486 and +RU-486) and 91% were common between the two groups. (E) Volcano plot illustrating significantly differentially abundant proteins. 40 proteins were found to be differentially expressed by using a cutoff on p value ≤ 0.05 and log_2_FC ≥ 1 (7 upregulated) and ≤ −1.0 (33 downregulated) proteins. (F) Twenty-five most significant biological processes that are downregulated upon overexpression of *chinmo* in adult neurons. See Supplementary figure for upregulated biological processes. (G, H) Overexpression of *chinmo* in the adult nervous system downregulates genes involved in fat metabolism. RT-PCR Quantitation of fold change in mRNA levels of genes (F) involved in fat metabolism in the head tissue (G) and decapitated body tissue (H) of *3X ElavGS > UAS chinmo* flies that were fed a solvent (light blue bars) or RU-486 (pink bars) diet for 10 days. (I) Knockdown of *FASN1* in the adult fat body reduces lifespan under AL (compare red solid and dotted lines) and DR (compare blue solid and blue dotted lines) conditions (See Supplementary table 5A for p values and median and maximum lifespan). (J) Knockdown of *FATP* in the adult fat body reduces lifespan under AL (compare red solid and dotted lines) and DR (compare blue solid and blue dotted lines) conditions (See Supplementary Table 5B for p values and median and maximum lifespan). (K) Overexpression of *Flag FATP* increases lifespan in flies that are fed a DR diet (Compare blue dotted line with blue solid line) (See Supplementary table 5B for p values and median and maximum lifespan).

### Overexpression of human primary *miR-125b-1* in the adult fat body extends lifespan

Given that DR regulated upregulation of *chinmo* was the likely cause of the decrease in lifespan extension upon DR, we tested whether increasing the levels of the brain enriched *miR-125* in adult fat body was able to mimic the beneficial effects of *chinmo_RNAi_*. Since, the human and fly processed miR-125 sequences are identical we generated transgenic fly lies that expressed the human primary *miR-125b-1* (*hsa pri miR-125b-1*) transcript (Figure 7A). Expression of *hsa primiR-125b-1* was induced in the adult fat body using the steroid inducible gene switch GAL4 driver *FB GS* (Figure 7B). Overexpression of *miR-125* in the adult fat body led to a 43% increase in maximum lifespan in AL diet and a 19.5% increase in median lifespan in DR diet (Figure 7C and supplementary Table 6). Consistent with the survival data, increasing *miR-125* levels in the fat tissue also led to increased stored triglyceride content in flies that were fed either AL or DR diet (Figure 7D). Moreover, *FB GS > UAS hsa pri miR-125b-1* flies that were fed a DR+ RU-486 diet displayed an increase in the diameter of the lipid droplets (Figure 7E). Finally, to test whether modulating *miR-125* levels led to changes in the expression of genes involved in fat metabolism, we examined the expression of *FASN1* and *FATP* in *FB GS > UAS hsa pri miR-125b-1* flies that were fed AL and DR in presence and absence of RU-486. Consistent with the increase in TAG levels, *FB GS > UAS hsa pri miR-125b-1* flies expressed higher levels of both *FASN1* and *FATP* in DR plus RU-486 conditions (Figure 7G-H). The levels of *FASN1* were also significantly higher in *FB GS > UAS hsa pri miR-125b-1* flies that were fed AL diet in presence of RU-486 (Figure 7H). These data confirmed that modulating the levels of *miR-125* and *chinmo* brought about an increase in lifespan by acting on the same downstream targets and that regulation of fat metabolism by *miR-125* is an evolutionary conserved mechanism.

**Figure 7.**
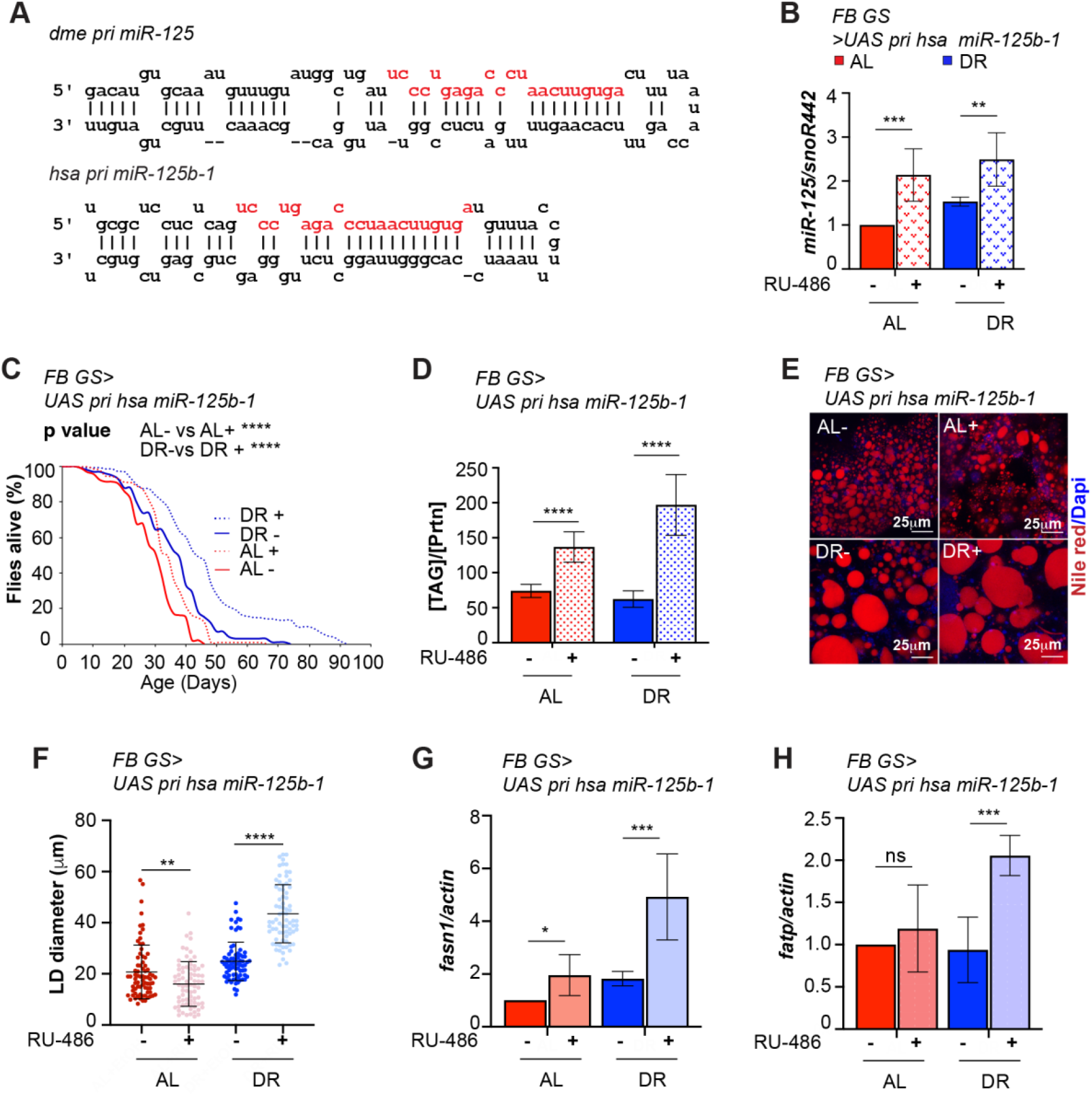
*miR-125* promotes longevity and fat metabolism. (A) Schematic of *Drosophila melanogaster* primary *miR-125* (*pri miR-125*) transcript and the *Homo sapiens* primary *miR-125b-1* (*pri hsa miR-125b-1*) indicating that the processed miRNA sequence in fruit flies and humans is identical. (B) Quantitative RT-PCR of *miR-125* from abdominal tissue of *FB GS > UAS chinmo* flies in presence of RU-486 (bars with red and blue pattern) or in absence of RU-486 (bars with solid red and blue color) flies that were fed *Ad libitum* (AL) (red) or DR diet (blue) for 20 days. Expression levels were normalized to SnoR442. Values are mean ± SD, n≥ 3. (C) Overexpression of *hsa pri miR-125* increases lifespan in flies that are fed an AL or DR diet (Compare red and blue dotted lines with red and blue solid line) (See Supplementary table 6 for p values and median and maximum lifespan). (D) *FB GS >UAS hsa pri miR-125b-1* flies store increased levels of triglycerides (TAG) under both AL and DR diets. Quantitation of TAG stored levels in -RU-486 and + RU-486 fed 20 d old *FBGS >UAS chinmo* flies. (E) Fat bodies/abdomens of female flies were dissected and stained for the content and diameter of lipid droplet (LD) (red are lipid droplets stained with Nile red and blue is Dapi). Scale bar, 25μm. (F) Quantitation of lipid droplet (LD) diameter in (E). Quantitation of 15 largest LDs in 5 samples per condition. Error bars represent ± SD and ** p <0.01. (G, H) Overexpression of *hsa pri miR-125* upregulates genes involved in fat metabolism under DR conditions. RT-PCR Quantitation of mRNA levels of *FASN1* (G) and FATP (H) in the fat tissue of *FB GS > UAS hsa pri mir-125* flies that were fed an AL (red and pink bars) or DR diet (dark blue and light blue) under uninduced (red and dark blue bars) and induced (pink and light blue bars) conditions for 20 days. Expression levels were normalized to *Actin5c*. Values are mean ± SD, n≥ 3.

## DISCUSSION

Regulatory RNAs are increasingly being recognized as key regulators of metabolic homeostasis.

In contrast to the protein machinery that represents only ~2% of the transcribed genome, the expansion of the noncoding transcriptome in higher eukaryotes reflects greater regulation of cellular processes through control of protein function(Consortium, 2012). Our previous analysis uncovered a role for *let-7* and *miR-125* in aging (Chawla et al., 2016). This work was followed up by another study that reported that increasing *Drosophila let-7* levels in the adult nervous system enhanced lifespan and altered metabolism(Gendron & Pletcher, 2017). In the current study, we have uncovered a new role for the evolutionary conserved *miR-125* in dietary restriction dependent extension of lifespan. We utilized hypomorphic and genetic loss of function mutants to examine the contribution of *miR-100, let-7* and *miR-125* in the DR pathway. *let-7-C_hyp_* mutant strain expresses near wild type levels of *let-7-C* miRNAs (*miR-100, let-7* and *miR-125*) during development but display an age-related decline in the levels of these miRNAs during adulthood (Chawla et al., 2016; Chawla & Sokol, 2012). Expression analysis of the *let-7-Complex* miRNAs in *let-7-C_hyp_* revealed that DR mediated upregulation of *let-7-C* miRNAs is required for lifespan extension by DR (Figure 1A, B). Lifespan analysis of *Δlet-7* and *ΔmiR-125* flies uncovered a role for these miRNAs in DR mediated lifespan extension. Thus, this is the first study that identifies miRNAs that are regulated by DR in the *Drosophila* model and demonstrates a role for these two miRNAs in DR mediated lifespan extension.

### Fat metabolism, DR and the role of miRNAs

Lipid metabolism plays an important role in the aging process and pharmacological, dietary and genetic interventions that extend lifespan often cause changes in lipid metabolism(Barzilai, Huffman, Muzumdar, & Bartke, 2012; Johnson & Stolzing, 2019). The adipose tissue has also been linked to metabolic dysfunction and age-related diseases such as heart attacks, stroke, hypertension, diabetes and cancer (Tchkonia et al., 2010). The fat body exerts some of its effects by storage and release of fat under different contexts. A non-autonomous regulatory role for this tissue in aging by regulation of the miRNA biogenesis machinery was reported in the mouse model. This study showed that aging associated decline in the levels of the miRNA biogenesis factor, Dicer results in the down regulation of a number of miRNAs including *miR-125b* in the adipose tissue (Mori et al., 2012). The authors further showed that knockout of Dicer specifically in the adipose tissue rendered the mice hypersensitive to oxidative stress. More importantly, this decline in multiple miRNAs in the adipose tissue was prevented by caloric restriction (CR). Though, this study correlated the age and CR mediated changes of *miR-125b* with its downstream target p53, the causal role of these changes on lifespan was not examined(Mori et al., 2012). Our genetic and molecular analysis in *Drosophila* independently identified *miR-125* as a downstream effector of DR and showed that upregulation of its human ortholog in the fat tissue was able to enhance lifespan.

### Chinmo as a nutrient dependent regulator of fat metabolism

Chinmo is a BTB (Bric-a-brac, Tramtrack, Broad complex) domain -Zinc finger (ZF) transcription factor that has been demonstrated to function as a repressor(Flaherty, Zavadil, Ekas, & Bach, 2009). This BTB-ZF protein has been shown to play a critical role in temporal fate specification of mushroom body neurons during development(Zhu et al., 2006). However, more recent studies have highlighted a new role for this protein in JAK/STAT signaling pathway that regulates stem-cell renewal in *Drosophila* (Flaherty et al., 2010). Interestingly, as a mediator JAK/STAT signaling, Chinmo exerts non-autonomous effects on renewal of germline stem cells in *Drosophila* testis(Flaherty et al., 2010). However, the molecular mechanism or the non-autonomous signal that mediates these effects are currently not known. Our study has uncovered a previously unknown functional role for this protein in fat metabolism. Proteomic analysis of flies that ectopically expressed *chinmo* in the adult neurons identified a number of downregulated biological processes including fat metabolism. In addition, we show that *chinmo* expression is regulated by nutrition. A significant increase in *chinmo* mRNA levels was observed in flies that were exposed to DR or starvation conditions, respectively (Figure 2G-I). These data reveal that Chinmo shares several attributes with the insulin/IGF signaling pathway--- it is a nutrient regulated factor that plays a critical role in temporal fate specification of neurons during development. Silencing of *chinmo* in adult brain by *miR-125* is required for normal aging(Chawla et al., 2016). More importantly, in this study we show that reducing *chinmo* in the adult fat body increases lifespan. Additionally, our results indicate that Chinmo may play a role in fat metabolism through its ability to circulate (Figure 4). Our expression analysis of Chinmo indicates that overexpression of *chinmo* in the adult nervous system results in its misexpression in several other peripheral tissue. Taken together these data identify Chinmo as a nutrient dependent regulator of aging.

### Role of miR-125 in aging and late onset diseases

We previously showed that *miR-125* plays a role in regulating lifespan and maintenance of neuronal integrity (Chawla et al., 2016). In this study we show that overexpression of the human *miR-125* in the adult fat tissue is sufficient to enhance lifespan. In addition to its role in modulating lifespan, *miR-125* has also been identified as a circulating diagnostic biomarker in Alzheimer’s disease and Type 2 Diabetes(Ortega et al., 2014; Tan et al., 2014; Villeneuve et al., 2010). These and several other studies, highlight the importance of this conserved small RNA as a prolongevity factor, biomarker and a disease modifier. However, the widespread expression of this miRNA in metazoans calls for a more detailed analysis of the tissue specific effects of *miR-125* before it can be considered as a relevant therapeutic target for treatment of age-related diseases.

Taken together, our analyses have identified *miR-125* as an effector of dietary restriction pathway. *miR-125* is regulated by dietary signals (DR) and represses *chinmo*, thus leading to de-repression of genes involved in fat metabolism in peripheral tissues, which in turn result in extension of lifespan (Figure 8). This functional analysis sets the stage for evaluation of *miR-125* and other conserved miRNAs as candidates for developing therapeutics that promote healthy aging and prevent/delay late onset diseases.

**Figure 8.**
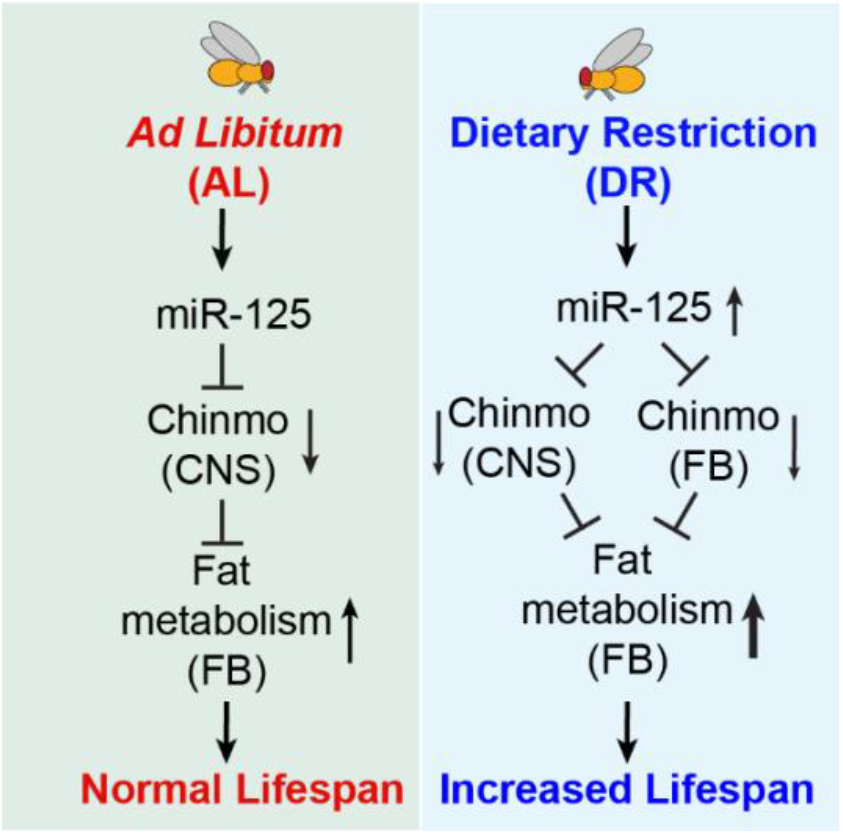
*miR-125* regulates DR-dependent lifespan extension by post-transcriptionally silencing chinmo. Proposed model summarizing the mechanism by which *miR-125* and *chinmo* regulate lifespan extension by dietary restriction.

## MATERIALS AND METHODS

### *Drosophila melanogaster* stocks and husbandry

Fly stocks were maintained in standard cornmeal/agar medium (composition provided in the next section) at 25°C with a 12h light: 12h dark cycle in 60% humidity. Fresh fly food was prepared every 3 days to prevent desiccation. For steroid mediated UAS-transgene control using the Gene-Switch driver, flies were fed a diet containing 200μM RU-486 (Mifepristone, Cayman Chemicals, Ann Arbor MI). Unless otherwise noted, adult female flies of indicated ages were used for experiments. All experiments were performed in the *Drosophila* chamber Model DR-36VL (Percival Scientific, Inc., IA, USA). Detailed genotypes of all strains as well as the sources of the genetic mutations and transgenes used in the study are listed Supplementary Tables S7 and S8 Tables, respectively. Transgenesis was performed by the Fly Facility at Bangalore Life Science Cluster (C, Bengaluru, India).

### Diets used in the study

*Cornmeal sugar medium (1L)*: Cornmeal (80g); Yeast extract (15g); Sucrose (25.85gm); Dextrose (51.65gm); 8g agar; methyl paraben (1gm) in 5ml Ethanol; 10mL of Acid mix (Propionic acid: Orthophosophoric acid).

*Ad libitum* (AL) diet (1L): Yeast Extract (5%); Corn meal (8.6%); Sucrose (5%); Agar (0.46%); 10ml of Acid Mix (Propionic acid: Orthophosophoric acid) (1%).

*Dietary restriction (DR) diet (1L)*: Yeast Extract (0.25%); Corn meal (8.6%); Sucrose (5%); Agar (0.46%); 10ml of Acid Mix (Propionic acid: Orthophosophoric acid) (1%).

*Starvation medium*: 1.5% agarose in Phosphate Buffered Saline (PBS).

### Survival analysis

Twenty female flies (1-3 day old) were transferred to each vial. Flies were transferred to fresh food every 3 days at which time dead flies were removed. Surviving flies were recorded every 2 days. The survival curves were plotted using Microsoft Excel. Statistical analysis was performed with the *Online Application for the Survival Analysis of lifespan assays* (OASIS) (Yang et al., 2011) and the p values were calculated using the log-rank (Mantel-cox) test. The number of flies used for each experiment are noted along with the median and maximum lifespans of the tested strains in Supplementary Tables S1, S2A-B, S3A-D, S4, S5A-C, S6. Experiments usually included two independent controls: *w_1118_* as well as a *let-7-C* mutant strain containing a fully rescuing transgene. The *w_1118_* survival curve was generated with flies that had been back crossed five times.

### Plasmid construction

The C-terminal Flag tagged *FATP* cDNA was sub-cloned as a XhoI-XbaI fragment into pUAST attB using the primers 211 and 209 by using plasmid SD05207 plasmid (DGRC). All PCRs were performed with High fidelity Phusion enzyme (Thermo Fisher Scientific, USA) and all clones were verified by sequencing. The *hsa mir-125b1* primary transcript was generated by annealing the oligo pair 1070/1071 and cloned into the XbaI site of pUAST attB.

### RNA isolation and quantitative real time PCR

Total RNA was extracted from whole fly or tissue samples using RNAiso Plus (Takara Bio, Inc). Animals/tissues were homogenized in 0.2 ml of RNAiso Plus with a micropestle (Tarsons) prior to extraction. The cDNA was generated by using a High capacity cDNA reverse transcription Kit (Thermo Fisher Scientific, MA, USA). In each reaction 0.5-1 μg was mixed with random hexamers, MgCl_2_, 10X RT Buffer, dNTPs, RNAse Inhibitor and MultiScribe Reverse transcriptase in a 10μl total volume. The cDNA synthesis was performed as per manufacturer’s protocol in a Bio-Rad C1000 Touch Thermal Cycler. The synthesized cDNA was diluted (1:10) and used as template for quantitative real-time PCR (qRT-PCR) using SYBR premix EX-Taq-plus (TaKara) and analyzed on QuantStudio 6 Real-Time PCR machine (Thermo Fisher Scientific, Waltham, Massachusetts, USA). The expression of the target genes was normalized to *actin-5C* or *rp49*. For expression analysis of miRNAs, Taqman miRNA assays (Thermo Fisher Scientific, Waltham, Massachusetts, USA) specific for the miRNA (dme miR-100, *dme-let-7* and *dme-miR-125*). Each cDNA sample was diluted 1:25 and real-time quantitative PCR (qPCR) was performed in duplicate using miRNA-specific primers/probe on a Quant 6 studio Real Time PCR System (Thermo Fisher Scientific, Waltham, Massachusetts, USA). For fold change analysis, individual values were normalized to 2S rRNA or Snu442 for Taqman miRNA assays and *rp49 or Actin 5c* levels for SYBR green assays. For qRT-PCR analysis, oligos 7, 8, 13, 14, 17, 18, 23, 24, 102a, 103a, 179, 180, 181, 182, 1183, 184, 185, 186, 187, 188, 189, 190, 191, 192, 193, 194, 3 and 4 listed in Table S9 were used.

### Measurement of protein levels

Protein levels were quantitated using the Bio-Rad protein assay dye reagent concentrate BIO-RAD, CA, USA) as per manufacturer’s instructions.

### Measurement of triglycerides (Pletcher et al.), glucose and glycogen levels

Triglyceride quantification was performed as described in (Tennessen, Barry, Cox, & Thummel, 2014).

### Immunostaining

Tissues were dissected in PBS and fixed for 30 minutes in 4% paraformaldehyde. For the staining of lipids, fruit flies were fed AL or DR diets for 20 days and fat bodies were dissected in 1X PBS. Tissues were fixed in 4% PFA for 30 minutes and 3 washes of 5 minutes each were given in 1X PBS+0.3% Triton. Tissues were stained for 1 hour with Nile Red Staining solution (1mg/mL in Acetone, 1:250 PBST) and DAPI (1:100 PBST). Samples were washed three times with PBST for 5 minutes and samples mounted. Immunofluorescence was performed as described previously (Chawla & Sokol, 2012; Wu et al., 2012). Primary antibodies included rat anti-Chinmo (Wu et al., 2012), rabbit anti-Woc (Raffa, Cenci, Siriaco, Goldberg, & Gatti, 2005) (gift from Maurizio Gatti 1:1000). Slides were analyzed under Leica SP8 Confocal Microscope and lipid droplet size was measured using Fiji software and graphs were plotted with GraphPad Prism v8. Confocal stacks were merged using Leica LAS software.

### Proteomics analysis

Age-matched mated female flies were fed + or + RU-486 food for 5 or 10 days. Protein lysate was prepared by freezing female flies (n=5 per replicate). Flies were homogenized in 100 μL of Ammonium bicarbonate buffer (100mM ammonium bicarbonate and 8M urea) after adding appropriate amount of 100X Protease inhibitor cocktail (Sigma). The samples were centrifuged for 15 minutes at maximum speed at 4°C in a microfuge and the supernatant was collected, quantitated and processed for Proteomic analysis.

#### Trypsin digestion and desalting

Proteins from control and 5 day treated samples were precipitated using 4 volume of chilled acetone. Precipitated proteins were washed thrice with chilled acetone and dried at room temperature. Precipitates were reconstituted in 100mM Ammonium bicarbonate and treated with 5mM DTT for 30 min at 55°C. Reduced disulphide bonds were alkylated using 10mM Iodoacetamide for 30 minute at 55°C. Proteins were digested using MS grade trypsin (Promega Trypsin Gold) in a ratio of 1:50 for 12 hrs. After digestion, peptides were dried in a SpeedVac vacuum dryer and reconstituted in 2% Acetonitrile (ACN) and 0.1% Formic acid (FA). The reconstituted peptides were cleaned and desalted using C18 cartridge (Oasis HLB 1cc). Samples were eluted in 0.1%FA and 50% ACN from the cartridge after desalting with 0.1% FA in water.

The eluted peptides were dried in a SpeedVac vacuum dryer and reconstituted in 15 μL of 2% ACN and 0.1%FA.

### LC-ESI MS analysis

Desalted peptides were loaded onto a CapTrap C18 trap cartridge Cap-Trap C18 trap cartridge (Michrom Bioresources, Auburn, CA, USA) and desalted for 10 minutes at the rate of 10 μl/ minute using 2% ACN and 0.05% trifluoroacetic acid (TFA) in water using Eksigent NanoLC 400. Labelled peptides were separated on Chromolith Caprod RP-18e HR capillary column (150 × 0.1 mm; Merck Millipore) using linear gradient of buffer B (98% ACN and 0.05% TFA in water) in buffer A (2% ACN and 0.05% TFA in water). Peptides were eluted at the rate of 300 nL/minute, which were directed to the 5600 TF for MS and MS/MS analysis. MS spectra were acquired from 350 Da to 1250 Da and MS/MS of the peptides were fragmented using IDA criteria. In brief, 25 most intense peaks were fragmented using collision induced dissociation (CID) with rolling collision energy in each cycle. The MS/MS spectra was acquired from 100 Da to 1600 Da.

### MS/MS Data Analysis

Protein search was performed using the database comprising of deduced amino acid sequences from *D. melanogaster*, available at UniProtKB website (https://www.uniprot.org/taxonomy/7227, proteome ID UP000000803). In order to avoid misidentifications, most common contaminants such as human keratin and porcine trypsin were also included in the database *.fasta file. The MS/MS peak list data files were analyzed by Mascot ion search engine version 2.3.02, with one fixed modification; carbamidomethylation of cysteine (monoisotopic mass of 57.0215 Da), two variable modifications; methionine oxidation (monoisotopic mass of 15.9949 Da), deamidation (monoisotopic mass of 0.984 Da) and a peptide and MS/MS fragment ion mass tolerance of 0.1 Da. Upto two missed cleavages were selected along with Mascot automatic decoy database search. The MASCOT DAT files from all fractions of each biological replica of 5 days control (1C1, 1C2 and 1C3) and treated (1E1, 1E2 and 1E3) samples were merged during the search and output file was analyzed using Scaffold Q+S version 4.9.0 (Proteome Software, Portland, OR, USA) to generate a full report of proteomic data, which is provided in Supplementary data (Excel sheets). Protein identification validation was performed by Scaffold parameters including Mascot ion scores of 30 or higher (for +2, +3 and +4 charges), a minimum of two identified peptides, 100 ppm of parent mass tolerance, 95% peptide identification probability and 95% protein identification probability, (using the Scaffold Local FDR algorithm), resulting in a 0.0% decoy FDR. Protein fold change values were calculated on the basis of Normalized Spectral Abundance Factor (NSAF), calculated for each protein. Proteins with fold change values less than or equal to 0.5 were down-regulated and the proteins with fold change values more than or equal to 2 were up-regulated.

Quantitative differences were statistically analysed by a t-test and volcano plot (Figure 6E), where differences with p values lower than 0.05 were considered statistically significant. A two-tailed t test with equal variance between samples was performed to calculate the p-value. In Figure 6E, INF (Infinity) values were changed to 4 and zero values to −4 (representing + and - or over and under expression). Identified proteins were categorized according to gene ontology terms using STRING, an online database of known and predicted protein-protein interactions. Validation of the identified proteins, spectral abundance values and comparative analysis between control and test samples was done using Scaffold. GO functional categories assigned to the identified proteins have been represented in (Figure 6F and Supplementary Excel sheets).

### Statistical analysis

Data representation and statistical analysis were performed using GraphPad Prism 8 software and/or Microsoft Excel. Survival curves were compared using log-rank tests, with Bonferroni corrections for p values where multiple comparisons were necessary. All survival and lifespan graphs show one representative experiment out of two or three independent repeats with 2-3 cohorts of 20 female flies per genotype. A two tailed t test was used to analyze data in Figure 1B, 2C-E, 2G-, 3F, H, 5A, 5C, 5E, 6A-C, 6G-H, 7B, 7D, 7F-H. Other details on statistical analysis can be found in Figure legends. Statistical significance was set at p<0.05. Asterisks indicate *p<0.05, **p< 0.01, ***p<0.001, ****p<0.0001, ns-non-significant, p≥0.05.

## ACKNOWLEDGEMENTS

This work was supported by the DBT/Wellcome Trust India Alliance Fellowship/Grant [grant number IA/I(S)/17/1/503085 awarded to GC. The proteomics analysis was performed at the Mass Spectrometry Facility of the Advanced Technology Platform Centre (ATPC) (Grant No. BT.MED-II/ATPC/BSC/01/2010). The authors acknowledge the advice and inputs of Dr. Nirpendra Singh for the proteomics analysis. Fly Facility at Bangalore Life Science Cluster is acknowledged for microinjection of the transgenic line constructs. The author’s than Dr. Arthur Luhur for microinjection of the *hsa miR-125b-1* construct. The authors thank Drs. Scott Pletcher and David Walker and Bloomington *Drosophila* Stock Center (NIH P40OD018537) for fly stocks, *Drosophila* Genomics Resource Center (NIH 2P40OD010949) for plasmids and Dr. M. Gatti for Woc antibody.

## COMPETING INTERESTS

No competing interests declared.

## Notes

### Competing Interest Statement

The authors have declared no competing interest.

