## Supplementary Data for "*miR-125-chinmo* pathway regulates dietary restriction dependent enhancement of lifespan in *Drosophila*"

### Supplementary Figures

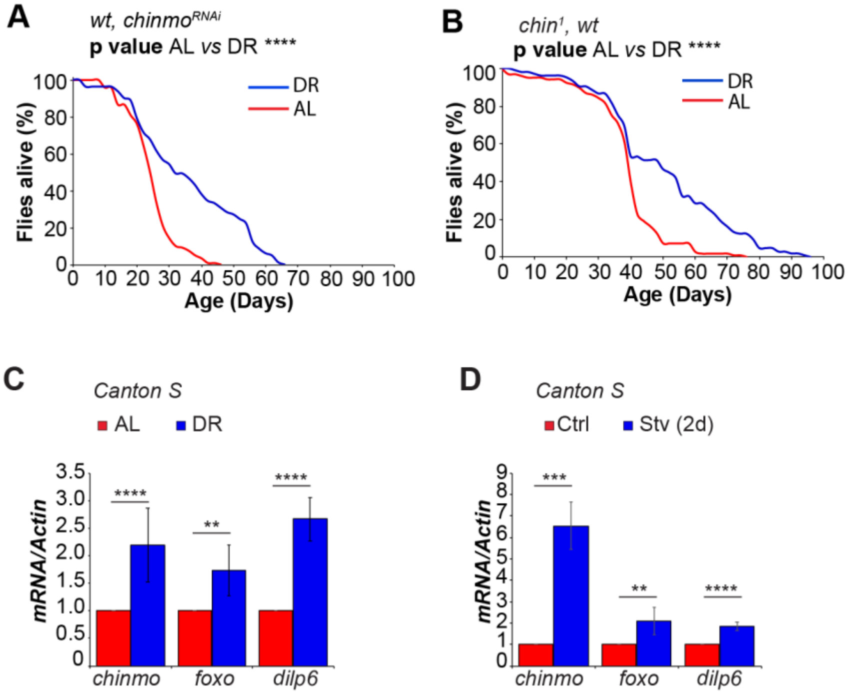

**Supplementary Figure 1. Reducing *chinmo* levels in the *let-7-CΔ* rescue line does not decrease the DR dependent life span extension.** (A, B) *Wild type (Rescue), UAS* $chinmo_{RNAi}$ and *wild type (Rescue),* $chin_1$ flies show a significant DR dependent increase in lifespan. P value < $10_{-10}$. For statistical analysis for comparison of survival curves, p values were calculated with log rank test. p value * <0.05. See Supplementary Table 2A, 2B for median and maximum lifespans. (C, D) DR and starvation induces expression of *chinmo* in wild type (*Canton S*) flies. (C) RT-PCR quantitation of fold change in *chinmo*, *foxo* and *dilp6* mRNA levels in *Canton S* flies that were fed an AL (Red bars) or DR (Blue bars) for 30 days. (D) RT-PCR Quantitation of fold change in *chinmo*, *foxo* and *dilp6* mRNA levels in *Canton S* flies that were fed a normal diet (Red bars) or starved (Blue bars) for 2 days. Expression levels were normalized to *Actin5c*. Values are mean ± SD, n≥ 3.

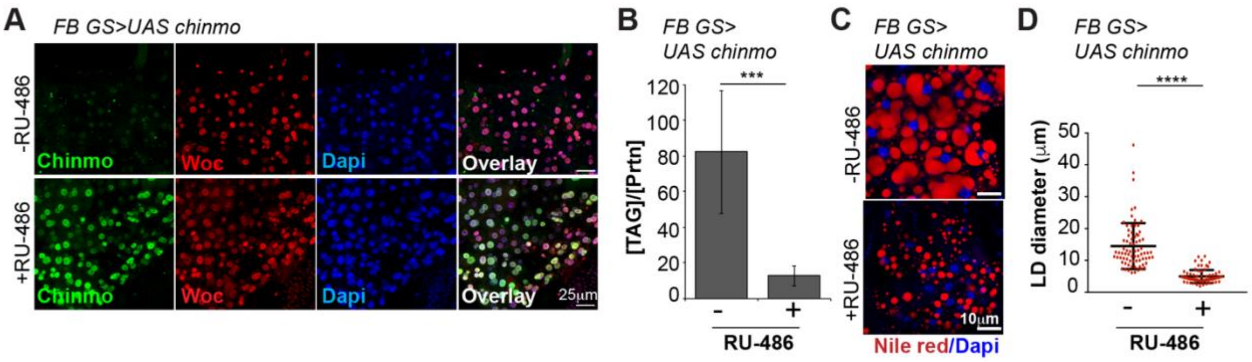

**Supplementary Figure 2. Ectopic expression of *chinmo* in the adult fat tissue reduces fat metabolism.** *UAS chinmo* was expressed in adult fat body using the drug (RU-486) inducible gene switch Gal4 driver. (A) Female flies that were fed an RU-486 supplemented diet for 5 days displayed increased levels of Chinmo in fat body cells, as detected by Chinmo (green), Woc (red) and Dapi (blue) staining of dissected fat body. (B) Quantitation of stored triglyceride (TAG) levels in -RU-486 and + RU-486 fed 20 d old *FBGS >UAS chinmo* flies. (C) Fat bodies/abdomens of female flies were dissected and stained for the content and diameter of lipid droplet (LD) (red are lipid droplets stained with Nile red and blue is Dapi). Scale bar, 25µm. (H) Quantitation of lipid droplet (LD) diameter in (C). Quantitation of 15 largest LDs in 5 samples per condition. Error bars represent ± SD and **** p <0.0001.

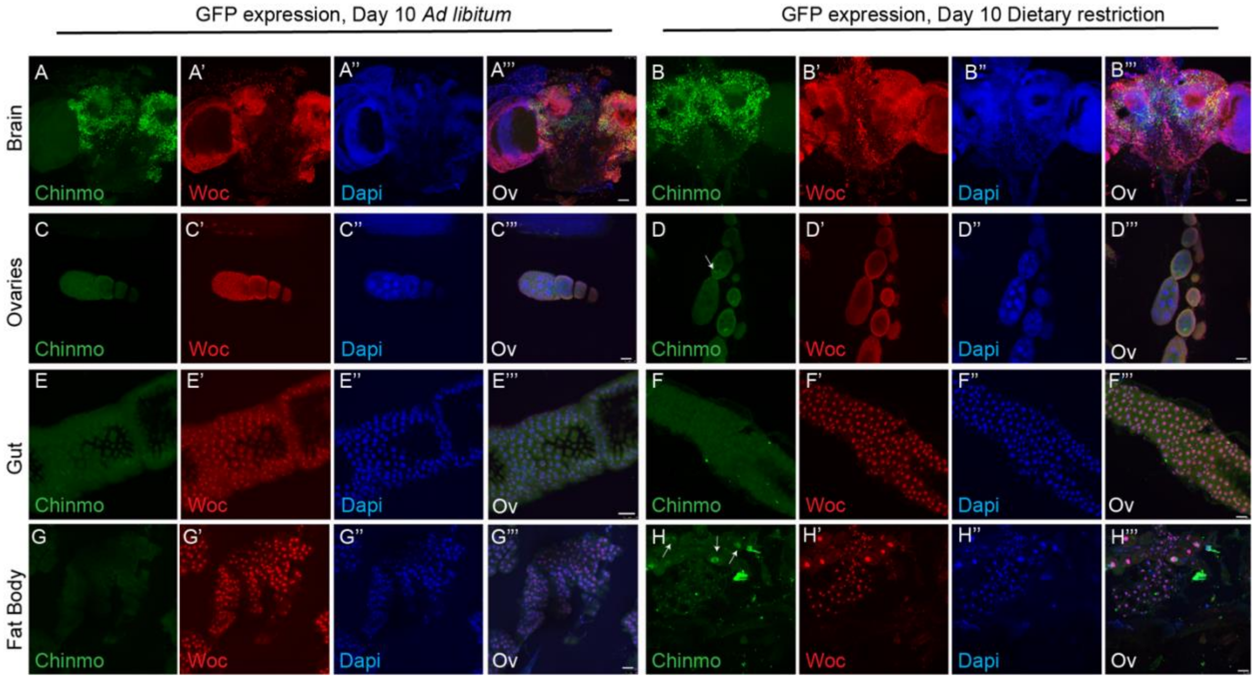

**Supplementary Figure 3. *miR-125* predominantly silences Chinmo in the adult brain.** Confocal images of 10d old adult brains (A-B), ovaries (C, D), gut (E, F) and fat body (G, H) of *miR-125* mutant flies that were fed an AL or DR diet for 10 days. The genotype of the flies in A-H is *let-7-C$_{GKI}$* / *let-7-C$_{KO2}$, P{neoFRT}40A; {v+, let-7-C $_{\Delta miR-125}$} attP2* / +. The dissected tissues were immunostained for Chinmo (green), Woc (red) and Dapi (blue). The intensity of Chinmo immunostaining is increased in ΔmiR-125 brains in both AL and DR diet (A-A''' and B-B'''). The ovaries, gut and fat body do not show any green signal in miR-125 mutants under AL conditions, indicating the chinmo is predominantly silenced in the brain by miR-125 in wild type flies. However, immunostaining of chinmo indicates some green signal in ovaries (D-D''') and fat body (H-H'''), indicating that the mRNA/protein of Chinmo is upregulated in these tissues under DR conditions. Scale bar, 25μm.

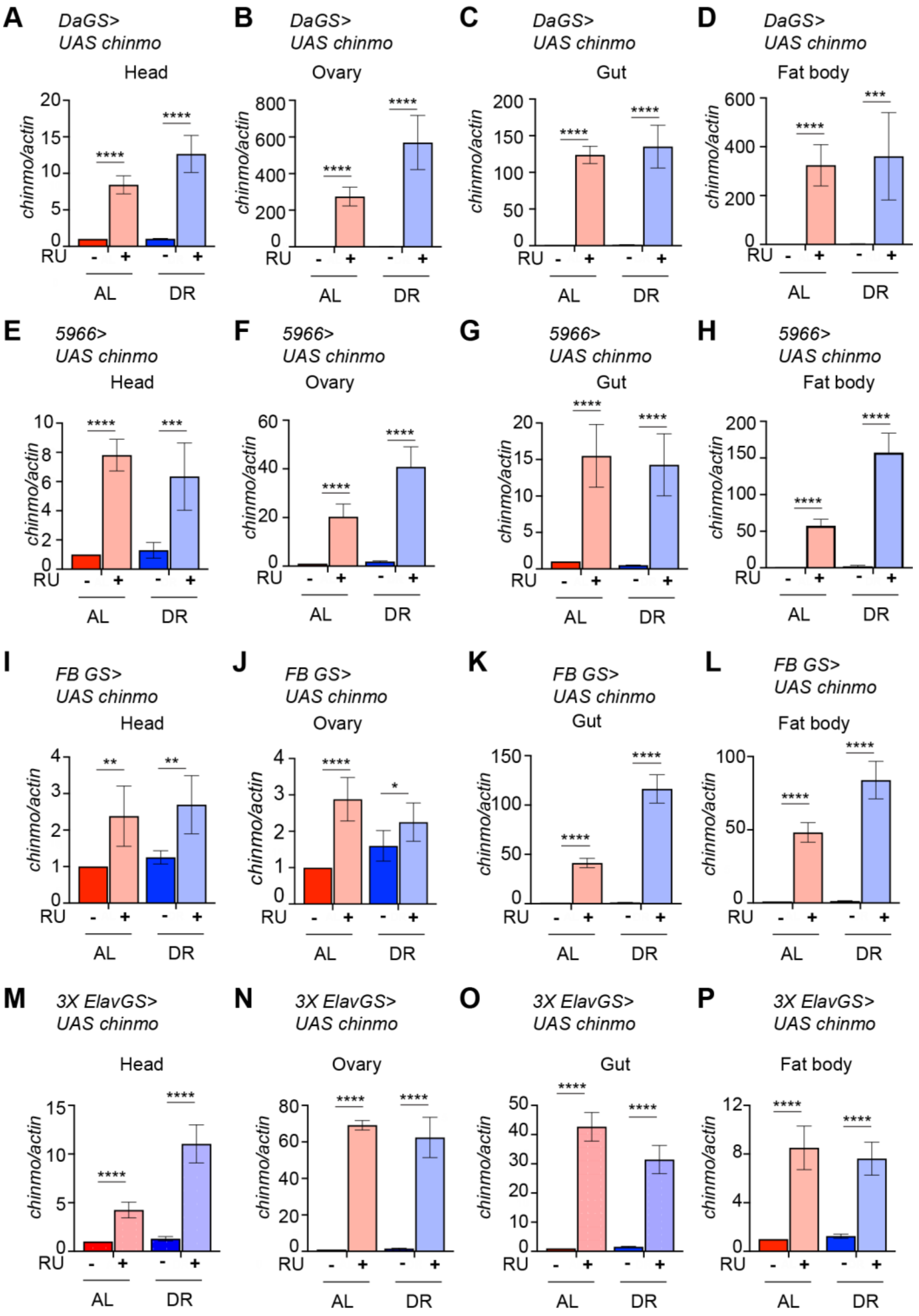

**Supplementary Figure 4. *Chinmo* mRNA is expressed non-autonomously upon overexpression.** *UAS chinmo* expression was driven in adult tissues in a drug inducible manner with a ubiquitously expressing *Da-GS* (A-D), *5966* (gut specific-GS) (E-H), fat body specific (*FB-GS*) (I-L) or neuronal (*3X*

*Elav-GS*) gene switch GAL4 drivers (M-P). (A-P) Quantitative RT-PCR of RNA extracted from dissected tissues (head, ovary, gut and fat body) from flies that were fed an AL (Red and pink bars) and DR (Dark Blue and Light blue bars) under uninduced (Red and Dark blue bars) and steroid induced (Pink and Light blue bar) conditions. Expression levels were normalized to *Actin5c*. Values are mean ± SD, n≥ 3.

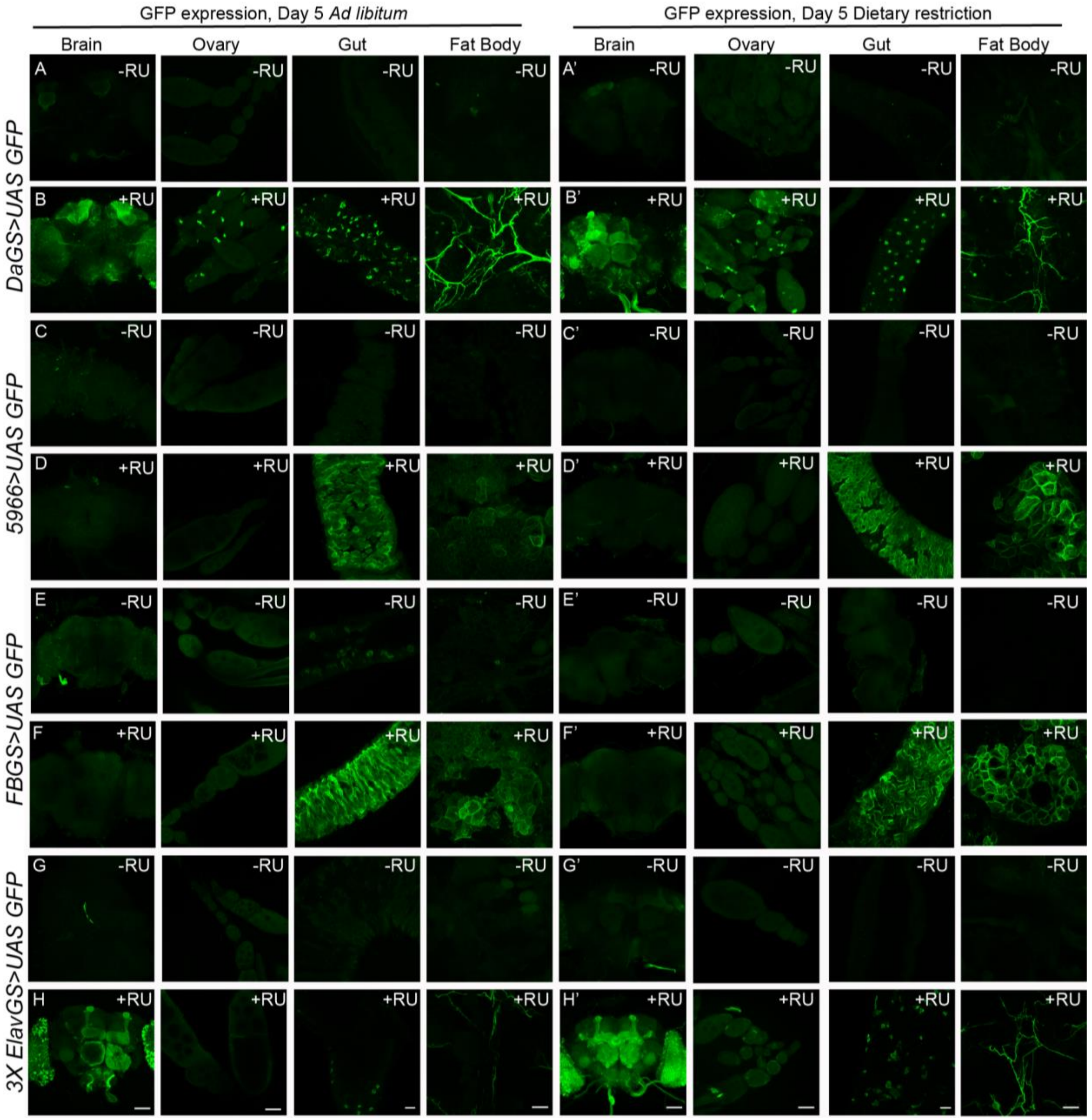

**Supplementary Figure 5. Verification of the specificity of the Gene switch Gal4 drivers.** *UAS GFP* was expressed in a steroid inducible manner with *Da-GS* (Supplementary for Figure 5A, 5A'-5B, 5B'), *5966* (gut specific-GS) (Supplementary for Figure 5C, 5C'-5D, 5D'), fat body specific (*FB-GS*) (Supplementary for Figure 5E, 5E'-5F, 5F') or neuronal (*3X Elav-GS*) gene switch GAL4 drivers (Supplementary for Figure 5G,5G'-5H, 5H'). (A-H') Confocal images of dissected *Drosophila* tissues (adult brain, ovaries, gut and fat body) from fruit flies that were fed an AL diet (5A-H) or DR diet (5A'-H') under uninduced (5A, 5C, 5E, 5G,

5A’,5C’, 5E’, 5G’) and steroid-induced (F, H, J, L, F’, H’, J’, L’) conditions. Scale bar, 50μm (brain, ovaries and fat body). Scale bar, 25μm (gut).

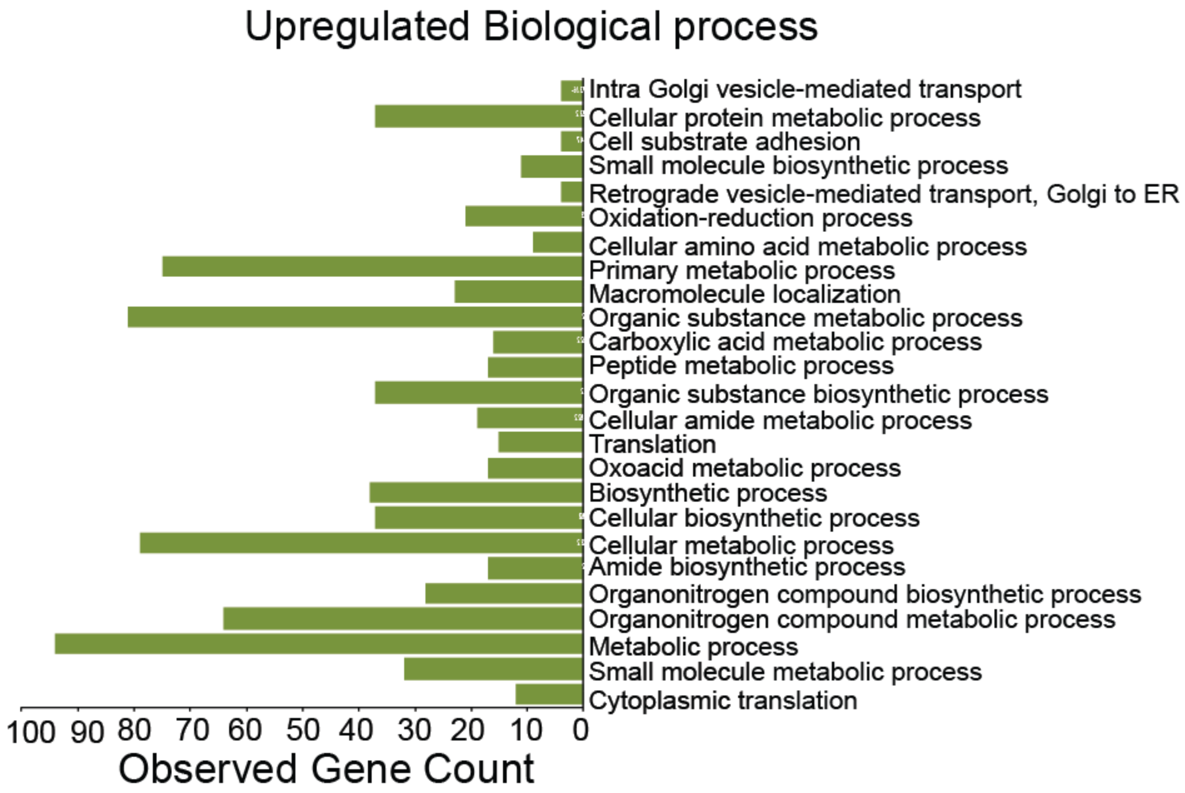

**Supplementary Figure 6. Over expression of *chinmo* in adult neurons upregulates cytoplasmic processes.** Twenty-five most significant biological processes that are upregulated upon overexpression of *chinmo* in adult neurons.

**Supplementary Table 1A.** Lifespan analysis of wild type and *let-7-Complex*$_{hyp}$ mutant line.

| | Lifespan (Days) | | p value** |
|---|---|---|---|
| #Experiment 1 | Maximum (Number of flies) | Median | |
| *w*$_{1118}$ *AL* | 50(99) | 28 | <1 x10$_{-10}$ |
| *w*$_{1118}$ *DR* | 88(98) | 58 | |
| *let-7-C*$_{hyp}$ *AL* | 40(123) | 20 | 0.2219 |
| *let-7-C*$_{hyp}$ *DR* | 40(125) | 20 | |
| Experiment 2 | | | |
| *w*$_{1118}$ *AL* | 50(153) | 24 | <1 x10$_{-10}$ |
| *w*$_{1118}$ *DR* | 60(125) | 32 | |
| *let-7-C*$_{hyp}$ *AL* | 42(138) | 22 | 0.0532 |
| *let-7-C*$_{hyp}$ *DR* | 44(164) | 24 | |
| Experiment 3 | | | |
| *w*$_{1118}$ *AL* | 54(186) | 22 | <1 x10$_{-10}$ |
| *w*$_{1118}$ *DR* | 82(162) | 40 | |
| *let-7-C*$_{hyp}$ *AL* | 36(196) | 20 | 0.1935 |
| *let-7-C*$_{hyp}$ *DR* | 42(165) | 18 | |

#Experiment 1 is represented in Figure 1; ** p value calculated by log rank test.

**Supplementary Table 1B.** Lifespan analysis of rescue, *ΔmiR-100*, *Δlet-7* and *ΔmiR-125* mutant lines.

| | **Lifespan (Days)** | | **p value****|
|---|---|---|---|
| *Experiment 1 | Maximum (Number of flies) | Median | |
| *Rescue AL* | 68(160) | 38 | $<1\ x10^{-10}$ |
| *Rescue DR* | 96(154) | 56 | |
| ΔmiR-100 *AL* | 56(199) | 32 | $<1\ x10^{-10}$ |
| ΔmiR-100 *DR* | 76(185) | 38 | |
| Δlet-7 *AL* | 40(86) | 20 | 0.0399 |
| Δlet-7 *DR* | 40(113) | 24 | |
| ΔmiR-125 *AL* | 48(146) | 28 | 0.281 |
| ΔmiR-125 *DR* | 52(139) | 28 | |
| Experiment 2 | | | |
| *Rescue AL* | 72(186) | 36 | $<1\ x10^{-10}$ |
| *Rescue DR* | 96(147) | 44 | |
| ΔmiR-100 *AL* | 56(196) | 28 | $<1\ x10^{-10}$ |
| ΔmiR-100 *DR* | 82(211) | 46 | |
| Δlet-7 *AL* | 42(86) | 18 | 0.0355 |
| Δlet-7 *DR* | 40(85) | 24 | |
| ΔmiR-125 *AL* | 48(144) | 30 | 0.0384 |
| ΔmiR-125 *DR* | 52(122) | 32 | |

*Experiment 1 is represented in Figure 1; **p value calculated by log rank test.

**Supplementary Table 2A.** Lifespan analysis of *rescue, chinmo*$_{RNAi}$ and *∆miR-125, chinmo*$_{RNAi}$ strains.

| | **Lifespan (Days)** | | **p value****|
|---|---|---|---|
| *Experiment 1 | Maximum (Number of flies) | Median | |
| *Rescue, chinmo*$_{RNAi}$ *AL* | 46(120) | 24 | 1.10E-07 |
| *Rescue, chinmo*$_{RNAi}$ *DR* | 62(81) | 32 | |
| ∆miR-125, *chinmo*$_{RNAi}$ *AL* | 36(86) | 24 | 3.50E-08 |
| ∆miR-125, *chinmo*$_{RNAi}$ *DR* | 50(101) | 32 | |
| Experiment 2 | | | |
| *Rescue, chinmo*$_{RNAi}$ *AL* | 48(145) | 28 | 0.0014 |
| *Rescue, chinmo*$_{RNAi}$ *DR* | 70(163) | 30 | |
| ∆miR-125, *chinmo*$_{RNAi}$ *AL* | 32(105) | 22 | 0.00E+00 |
| ∆miR-125, *chinmo*$_{RNAi}$ *DR* | 40(98) | 28 | |

*Experiment 1 is represented in Figure 2 and Supplementary Fig 2; **p value calculated by log rank test.

**Supplementary Table 2B.** Lifespan analysis of *rescue, chinmo*$_1$ and *ΔmiR-125, chinmo*$_1$ strains.

| | **Lifespan (Days)** | | **p value**** |
|---|---|---|---|
| *Experiment 1 | Maximum (Number of flies) | Median | |
| *Rescue, chinmo*$_1$ *AL* | 76(110) | 40 | 0.00E+00 |
| *Rescue, chinmo*$_1$ *DR* | 96(105) | 50 | |
| ΔmiR-125, *chinmo*$_1$ *AL* | 64(114) | 34 | 0.00E+00 |
| ΔmiR-125, *chinmo*$_1$ *DR* | 98(98) | 58 | |
| Experiment 2 | | | |
| *Rescue, chinmo*$_1$ *AL* | 84(128) | 38 | 2.3 x10$^{-9}$ |
| *Rescue, chinmo*$_1$ *DR* | 98(147) | 46 | |
| ΔmiR-125, *chinmo*$_1$ *AL* | 56(85) | 28 | 0.00E+00 |
| ΔmiR-125, *chinmo*$_1$ *DR* | 86(97) | 54 | |
| Experiment 3 | | | |
| *Rescue, chinmo*$_1$ *AL* | 78(127) | 32 | 0.0074 |
| *Rescue, chinmo*$_1$ *DR* | 96(182) | 34 | |
| ΔmiR-125, *chinmo*$_1$ *AL* | 60(95) | 24 | 0.00E+00 |
| ΔmiR-125, *chinmo*$_1$ *DR* | 94(106) | 38 | |

*Experiment 1 is represented in Figure 2 and Supplementary Fig 2; **p value calculated by log rank test.

**Supplementary Table 3A.** Lifespan analysis of *DaGS> UAS chinmo* strain.

| | **Lifespan (Days)** | | **p value**** |
|---|---|---|---|
| #Experiment 1 | Maximum (Number of flies) | Median | |
| *DaGS> UAS chinmo AL -RU* | 48(105) | 32 | 0.00E+00 |
| *DaGS> UAS chinmo DR -RU* | 80(108) | 52 | |
| *DaGS> UAS chinmo AL +RU* | 12(124) | 6 | 0.3895 |
| *DaGS> UAS chinmo DR +RU* | 12(132) | 6 | |
| Experiment 2 | | | |
| *DaGS> UAS chinmo AL -RU* | 46(158) | 30 | 0.00E+00 |
| *DaGS> UAS chinmo DR -RU* | 78(99) | 46 | |
| *DaGS> UAS chinmo AL +RU* | 12(118) | 6 | 0.0001 |
| *DaGS> UAS chinmo DR +RU* | 12(111) | 8 | |
| Experiment 3 | | | |
| *DaGS> UAS chinmo AL -RU* | 50(189) | 28 | 0.00E+00 |
| *DaGS> UAS chinmo DR -RU* | 76(113) | 52 | |
| *DaGS> UAS chinmo AL +RU* | 12(139) | 6 | 0.0001 |
| *DaGS> UAS chinmo DR +RU* | 12(123) | 8 | |

#Experiment 1 is represented in Figure 3; ** p value calculated by log rank test.

**Supplementary Table 3B.** Lifespan analysis of *5966> UAS chinmo* strain.

| | **Lifespan (Days)** | | **p value**** |
|---|---|---|---|
| #Experiment 1 | Maximum (Number of flies) | Median | |
| *5966> UAS chinmo AL -RU* | 56(99) | 24 | $5.2 \times 10^{-8}$ |
| *5966> UAS chinmo DR -RU* | 82(108) | 34 | |
| *5966> UAS chinmo AL +RU* | 40(97) | 12 | 0.0035 |
| *5966> UAS chinmo DR +RU* | 52(85) | 12 | |
| Experiment 2 | | | |
| *5966> UAS chinmo AL -RU* | 60(96) | 26 | $6 \times 10^{-4}$ |
| *5966> UAS chinmo DR -RU* | 74(97) | 34 | |
| *5966> UAS chinmo AL +RU* | 38(103) | 12 | 0.2405 |
| *5966> UAS chinmo DR +RU* | 58(93) | 12 | |
| Experiment 3 | | | |
| *5966> UAS chinmo AL -RU* | 78(89) | 26 | $2.3 \times 10^{-5}$ |
| *5966> UAS chinmo DR -RU* | 88(88) | 32 | |
| *5966> UAS chinmo AL +RU* | 44(93) | 10 | 0.0114 |
| *5966> UAS chinmo DR +RU* | 56(99) | 12 | |

#Experiment 1 is represented in Figure 3; ** p value calculated by log rank test.

**Supplementary Table 3C.** Lifespan analysis of *FBGS> UAS chinmo*

| | Lifespan (Days) | | p value** |
|---|---|---|---|
| #Experiment 1 | Maximum (Number of flies) | Median | |
| *FBGS> UAS chinmo AL -RU* | 64(81) | 34 | 0.00E+00 |
| *FBGS> UAS chinmo DR -RU* | 80(91) | 62 | |
| *FBGS> UAS chinmo AL +RU* | 32(106) | 18 | 0.3035 |
| *FBGS> UAS chinmo*$_i$ *DR +RU* | 38(104) | 20 | |
| Experiment 2 | | | |
| *FBGS> UAS chinmo AL -RU* | 38(105) | 22 | 1.30E-06 |
| *FBGS> UAS chinmo DR -RU* | 50(105) | 30 | |
| *FBGS> UAS chinmo AL +RU* | 36(91) | 16 | 0.0011 |
| *FBGS> UAS chinmo DR +RU* | 34(98) | 20 | |
| Experiment 3 | | | |
| *FBGS> UAS chinmo AL -RU* | 38(105) | 22 | 0.00E+00 |
| *FBGS> UAS chinmo DR -RU* | 72(96) | 40 | |
| *FBGS> UAS chinmo AL +RU* | 24(107) | 12 | 0.0053 |
| *FBGS> UAS chinmo DR +RU* | 42(105) | 14 | |

#Experiment 1 is represented in Figure 4; ** p value calculated by log rank test.

**Supplementary Table 3D.** Lifespan analysis of *3X ElavGS > UAS chinmo.*

| | **Lifespan (Days)** | | **p value**** |
|---|---|---|---|
| #Experiment 1 | Maximum (Number of flies) | Median | |
| *3X ElavGS> UAS chinmo AL -RU* | 48(97) | 22 | 0.0e+00 |
| *3X ElavGS > UAS chinmo DR -RU* | 72(80) | 56 | |
| *3X ElavGS > UAS chinmo AL +RU* | 48(95) | 14 | 0.3299 |
| *3X ElavGS > UAS chinmo DR +RU* | 58(97) | 16 | |
| Experiment 2 | | | |
| *3X ElavGS > UAS chinmo AL -RU* | 56(98) | 32 | 0.0e+00 |
| *3X ElavGS > UAS chinmo DR -RU* | 88(99) | 42 | |
| *3X ElavGS > UAS chinmo AL +RU* | 50(100) | 10 | 0.0066 |
| *3X ElavGS > UAS chinmo DR +RU* | 52(95) | 16 | |
| Experiment 3 | | | |
| *3X ElavGS > UAS chinmo AL -RU* | 58(145) | 28 | 0.0e+00 |
| *3X ElavGS > UAS chinmo DR -RU* | 80(88) | 42 | |
| *3X ElavGS > UAS chinmo AL +RU* | 56(131) | 14 | 0.0898 |
| *3X ElavGS > UAS chinmo DR +RU* | 56(82) | 18 | |

#Experiment 1 is represented in Figure 3; ** p value calculated by log rank test.

**Supplementary Table 4.** Lifespan analysis of *FBGS> UAS chinmo*$_{RNAi}$ strain.

| | Lifespan (Days) | | p value** |
|---|---|---|---|
| #Experiment 1 | Maximum (Number of flies) | Median | |
| *FBGS> UAS chinmo*$_{RNAi}$ *DR-RU* | 92(138) | 62 | 0.0408 |
| *FBGS> UAS chinmo*$_{RNAi}$ *DR +RU* | 102(88) | 60 | |
| *FBGS> UAS chinmo*$_{RNAi}$ *AL-RU* | 72(97) | 42 | 5.4 x10$^{-6}$ |
| *FBGS> UAS chimo*$_{RNAi}$ *AL+RU* | 80(98) | 50 | |
| Experiment 2 | | | |
| *FBGS> UAS chinmo*$_{RNAi}$ *DR-RU* | 96(100) | 38 | 2.4 x10$^{-8}$ |
| *FBGS> UAS chinmo*$_{RNAi}$ *DR +RU* | 100(102) | 82 | |
| *FBGS> UAS chinmo*$_{RNAi}$ *AL-RU* | 48(82) | 30 | 0.00E+00 |
| *FBGS> UAS chinmo*$_{RNAi}$ *AL+RU* | 70(92) | 42 | |
| Experiment 3 | | | |
| *FBGS> UAS chinmo*$_{RNAi}$ *DR-RU* | 64(105) | 42 | 0.00E+00 |
| *FBGS> UAS chinmo*$_{RNAi}$ *DR +RU* | 90(113) | 52 | |
| *FBGS> UAS chinmo*$_{RNAi}$ *AL-RU* | 58(100) | 30 | 0.0224 |
| *FBGS> UAS chinmo*$_{RNAi}$ *AL+RU* | 66(86) | 34 | |

#Experiment 1 is represented in Figure 3; ** p value calculated by log rank test.

**Supplementary Table 5A.** Lifespan analysis of *FBGS> UAS* $FASN1_{RNAi}$

| | **Lifespan (Days)** | | **p value**** |
|---|---|---|---|
| #Experiment 1 | Maximum (Number of flies) | Median | |
| *FBGS> UAS* $FASN1_{RNAi}$ *AL -RU* | 54(148) | 34 | 2.00E-05 |
| *FBGS> UAS* $FASN1_{RNAi}$ *AL +RU* | 44(99) | 30 | |
| *FBGS> UAS* $FASN1_{RNAi}$ *DR -RU* | 66(80) | 44 | 0.00E+00 |
| *FBGS> UAS* $FASN1_{RNAi}$ *DR +RU* | 52(60) | 30 | |
| Experiment 3 | | | |
| *FBGS> UAS* $FASN1_{RNAi}$ *AL -RU* | 46(80) | 28 | 0.0013 |
| *FBGS> UAS* $FASN1_{RNAi}$ *AL +RU* | 36(64) | 28 | |
| *FBGS> UAS* $FASN1_{RNAi}$ *DR -RU* | 82(56) | 38 | 0.00E+00 |
| *FBGS> UAS* $FASN1_{RNAi}$ *DR +RU* | 66(67) | 42 | |

#Experiment 1 is represented in Figure 5; ** p value calculated by log rank test.

**Supplementary Table 5B.** Lifespan analysis of *FBGS> UAS FATP*$_{RNAi}$.

| | Lifespan (Days) | | p value** |
|---|---|---|---|
| #Experiment 1 | Maximum (Number of flies) | Median | |
| *FBGS> UAS FATP*$_{RNAi}$ *AL -RU* | 62(113) | 32 | 0.0001 |
| *FBGS> UAS FATP*$_{RNAi}$ *AL +RU* | 62(130) | 40 | |
| *FBGS> UAS FATP*$_{RNAi}$ *DR -RU* | 94(97) | 46 | 3.00E-05 |
| *FBGS> UAS FATP*$_{RNAi}$ *DR +RU* | 74(124) | 40 | |
| Experiment 2 | | | |
| *FBGS> UAS FATP*$_{RNAi}$ *AL -RU* | 62(95) | 40 | 0.0006 |
| *FBGS> UAS FATP*$_{RNAi}$ *AL +RU* | 62(146) | 34 | |
| *FBGS> UAS FATP*$_{RNAi}$ *DR -RU* | 96(79) | 54 | 0.0002 |
| *FBGS> UAS FATP*$_{RNAi}$ *DR +RU* | 86(100) | 46 | |

#Experiment 1 is represented in Figure 6; ** p value calculated by log rank test.

**Supplementary Table 5C.** Lifespan analysis of *FBGS> UAS FATP* flies.

| | **Lifespan (Days)** | | **p value**** |
|---|---|---|---|
| #Experiment 1 | Maximum (Number of flies) | Median | |
| *FBGS> UAS FATP AL -RU* | 42(56) | 30 | 0.6555 |
| *FBGS> UAS FATP AL +RU* | 44(79) | 28 | |
| *FBGS> UAS FATP DR -RU* | 60(80) | 42 | 0.00E+00 |
| *FBGS> UAS FATP DR +RU* | 70(104) | 46 | |
| Experiment 2 | | | |
| *FBGS> UAS FATP AL -RU* | 36(93) | 24 | 0.00E+00 |
| *FBGS> UAS FATP AL +RU* | 50(94) | 34 | |
| *FBGS> UAS FATP DR -RU* | 36(95) | 30 | 0.00E+00 |
| *FBGS> UAS FATP DR +RU* | 60(80) | 40 | |

#Experiment 1 is represented in Figure 6; ** p value calculated by log rank test.

**Supplementary Table S6.** Lifespan analysis of *FB GS> UAS pri hsa miR-125b-1 strain.*

| | Lifespan (Days) | | p value** |
|---|---|---|---|
| #Experiment 1 | Maximum (Number of flies) | Median | |
| *FB GS> UAS pri hsmiR-125 AL -RU* | 50(93) | 32 | 3.9 x10$^{-5}$ |
| *FB GS> UAS pri hsmiR-125 AL +RU* | 88(145) | 34 | |
| *FB GS> UAS pri hsmiR-125 DR -RU* | 74(99) | 38 | 1.4 x10$^{-7}$ |
| *FB GS> UAS pri hsmiR-125 DR +RU* | 92(135) | 44 | |
| Experiment 2 | | | |
| *FB GS> UAS pri hsmiR-125 AL -RU* | 54(106) | 32 | 0.0001 |
| *FB GS> UAS pri hsmiR-125 AL +RU* | 68(144) | 36 | |
| *FB GS> UAS pri hsmiR-125 DR -RU* | 64(108) | 44 | 1.4 x10$^{-5}$ |
| *FB GS> UAS pri hsmiR-125 DR +RU* | 92(134) | 48 | |
| Experiment 3 | | | |
| *FB GS> UAS pri hsmiR-125 AL -RU* | 44(89) | 30 | 1.4 x10$^{-5}$ |
| *FB GS> UAS pri hsmiR-125 AL +RU* | 68(85) | 34 | |
| *FB GS> UAS pri hsmiR-125 DR -RU* | 84(112) | 40 | 0.0003 |
| *FB GS> UAS pri hsmiR-125 DR +RU* | 96(120) | 48 | |

#Experiment 1 is represented in Figure 7; ** p value calculated by log rank test.

#### Supplementary Table 7. Genotypes used in this study.

| Fig. | Strain Name | Genotype |
|---|---|---|
| 1A-G | *Wt* | $w_{1118}$ |
| | *let-7-C*$_{hyp}$ | $w_{1118}$; *let-7-C*$_{GKI}$/ *let-7-C*$_{KO2}$, *P{neoFRT}40A ; P{w+, let-7-Cp*$_{3.3kb}$*::cDNA}VK00033 / {v+, let-7-C* $_{\Delta let\text{-}7\text{-}C\ miRNAs}$*}attP2* |
| | *Rescue (let-7-C*$_{null}$ *rescue)* | $w_{1118}$; *let-7-C*$_{GKI}$ / *let-7-C*$_{KO2}$, *P{neoFRT}40A ; {v+, let-7-C}attP2 / +* |
| | *ΔmiR-100* | $w_{1118}$; *let-7-C*$_{GKI}$ / *let-7-C*$_{KO2}$, *P{neoFRT}40A ; {v+, let-7-C* $_{\Delta miR\text{-}100}$*}attP2 / +* |
| | *Δlet-7* | $w_{1118}$; *let-7-C*$_{GKI}$ / *let-7-C*$_{KO2}$, *P{neoFRT}40A ; {v+, let-7-C* $_{\Delta let\text{-}7}$*}attP2 / +* |
| | *ΔmiR-125* | $w_{1118}$; *let-7-C*$_{GKI}$ / *let-7-C*$_{KO2}$, *P{neoFRT}40A ; {v+, let-7-C* $_{\Delta miR\text{-}125}$*}attP2 / +* |
| 2 | *chin*$_{1}$ *, wt* | $w_{1118}$; *let-7-C*$_{GKI}$ / *chinmo*$_{1}$, *let-7-C*$_{KO2}$, *P{neoFRT}40A ; {v+, let-7-C}attP2 / +* |
| | *chin*$_{1}$, *ΔmiR-125* | $w_{1118}$; *let-7-C*$_{GKI}$ / *chinmo*$_{1}$, *let-7-C*$_{KO2}$, *P{neoFRT}40A ; {v+, let-7-C* $_{\Delta miR\text{-}125}$*}attP2 / +* |
| | *chin*$_{RNAi}$, *ΔmiR-125* | $w_{1118}$; *let-7-C*$_{GKI}$ / *let-7-C*$_{KO2}$, *P{neoFRT}40A ; {v+, let-7-C* $_{\Delta miR\text{-}125}$*}attP2 / P{w+, UAS-chinmo*$_{RNAi\ 148}$*}VK00033* |
| | *Wt* | *Canton S* |
| | *Wt* | $w_{1118}$ |
| 3A-H and 4A-H' | *DaGS, UAS-chinmo* | *w[1118]; P{Da GS }/+; P{w+, UAS-chin::SV40}/+* |
| | *5966, UAS-chinmo* | *w[1118]; P{5966 GS }/+; P{w+, UAS-chin::SV40}/+* |
| | *FB-GS, UAS-chinmo* | *w[1118]; P{w[+mW.hs]=Switch1}106/+; P{w+, UAS-chin::SV40}/+* |
| | *3XelavGS, UAS-chinmo* | *P{elav-Switch.O}GS -1A / + ; P{elav-Switch.O}GS-3A, P{elav-Switch.O}GSG301 / P{w+, UAS-chin::SV40}/+* |
| 5A-E | *FB-GS, UAS Chinmo*$_{RNAi}$ | *w[1118]; P{w[+mW.hs]=Switch1}106/+; P{w+, UAS-chinmo*$_{RNAi\ 148}$*}VK00033* |
| 6A-K | *3X ElavGS, UAS chinmo* | *P{elav-Switch.O}GS -1A / + ; P{elav-Switch.O}GS-3A, P{elav-Switch.O}GSG301 / P{w+, UAS-chin::SV40}/+* |
| | *FB GS, UAS FASN*$_{RNAi}$ | *w[1118]; P{w[+mW.hs]=Switch1}106/+; P{y[+t7.7] v[+t1.8]=TRiP.HMS01524}attP2/+* |
| | *FB GS, UAS FATP*$_{RNAi}$ | *w[1118]; P{w[+mW.hs]=Switch1}106/+; P{y[+t7.7] v[+t1.8]=TRiP.HMC04206}attP2/+* |
| | *FB GS, UAS Flag FATP* | *w[1118]; P{w[+mW.hs]=Switch1}106/+;P{w+, UAS-Flag FATP} attP2 / +* |
| 7B-H | *FB GS, UAS has miR-125b-1* | *w[1118]; P{w[+mW.hs]=Switch1}106/+; P{v+, UAS-hsa miR-125b-1} VK00033/+* |

#### Supplementary Table 8. Sources of mutations and transgenes used in study

| Strain | Source |
|---|---|
| *let-7-C*$_{GKI}$ | Ref (Sokol et al., 2008) |
| *let-7-C*$_{KO2}$ | Ref (Wu et al., 2012) |
| *chinmo*$_{1}$ | Ref (Zhu et al., 2006) |
| *P{w+, UAS-Chin::SV40}* | Ref (Chawla et al., 2016) |
| *P{w+, UAS-chinmo*$_{RNAi\ 148}$*}VK00033* | Ref (Chawla et al., 2016) |
| *P{w+, UAS-hsa miR-125B-1} VK00033* | This study |
| *P{v+, let-7-C* $_{\Delta miR\text{-}100}$*}attP2* | Ref (Chawla et al., 2016) |
| *P{v+, let-7-C* $_{\Delta let\text{-}7}$*}attP2* | Ref (Chawla et al., 2016) |
| *P{v+, let-7-C*$_{\Delta miR\text{-}125}$*}attP2* | Ref (Chawla et al., 2016) |
| *P{w+, let-7-Cp*$_{3.3kb}$*::cDNA}VK00033* | Ref (Chawla et al., 2016) |
| *3XelavGS* | Kind gift from Scott Pletcher |
| *P{w+, UAS-FASN1*$_{RNAi\text{-}1}$*}attP2* | Bloomington Stock 35775 |
| *P{w+, UAS-FATP*$_{RNAi}$*}attP2* | Bloomington Stock 55919 |
| *P{w+, UAS-Flag FATP::SV40}* | This study |
| *FB-GS* | Bloomington Stock 8151 |
| *5966 (Gut specific GS)* | Kind gift from David Walker |
| *Da GS* | Kind gift from David Walker |
| *UAS GFP* | Bloomington Stock 32198 |

**Supplementary Table 9. Primers used in this study.**

| No. | Name | Sequence |
|---|---|---|
| 7 | Chinmo QPCR For | AGTTCTGCCTCAAATGGAACAG |
| 8 | Chinmo QPCR Rev | CGCAGGATAATATGACATCGGC |
| 13 | Foxo QPCR-1 For | TCGCCGAACTCAGTAACCAC |
| 14 | Foxo QPCR-1 Rev | CACCTCCAGGCATTGTCCTT |
| 17 | Dilp 6 QPCR For | CGATGTATTTCCCAACAGTTTCG |
| 18 | Dilp 6 QPCR Rev | AAATCGGTTACGTTCTGCAAGTC |
| 19 | Dilp 2 QPCR For | AGCAAGCCTTTGTCCTTCATCTC |
| 20 | Dilp 2 QPCR Rev | ACACCATACTCAGCACCTCGTTG |
| 23 | act-5c QPCR For | CACACCAAATCTTACAAAATGTGT |
| 24 | act-5c QPCR Rev | AATCCGGCCTTGCACATG |
| 102a | FASN 1 RT PCR For | GTGCGTCCTATCAGCTACCC |
| 103a | FASN 1 RT PCR Rev | GTCTGCCAAGCCAGAGTCAT |
| 1004 | CHUTRSHRNA.3T(NS) | ctagcagtccaactgaatttcaattgtgatagttatattcaagcatatcacaattgaaattcagttgggcg |
| 1005 | CHUTRSHRNA.3B(NS) | aattcgcccaactgaatttcaattgtgatatgcttgaatataactatcacaattgaaattcagttggactg |
| 179 | FATP RT-PCR For | GCGGTTATCTCTCACTCCC |
| 180 | FATP RT-PCR Rev | CGCAGTCGGCGAAATAGTTG |
| 181 | CG2107 RT PCR For | CCACACGGGACTTCTGTGAA |
| 182 | CG2107 RT PCR Rev | ATGCGCTTGTAAGCCTCAGT |
| 183 | CG5009 RT PCR For | AGACCAGGGCTGACTACGAT |
| 184 | CG5009 RT PCR Rev | CATGGGACGGTGAGTATCGG |
| 185 | CG8778 RT PCR For | AGAGCGTCTGGTTTAGCTCG |
| 186 | CG8778 RT PCR Rev | CCTCGGGAGTCATGCCTTTT |
| 187 | CG9527 RT PCR For | ACTTCCGTAGCGGACCTTTG |
| 188 | CG9527 RT PCR Rev | GCAGAAGATGTGGGGTTCCA |
| 189 | CG9577 RT PCR For | GACTGGCCACTAATCCCGAC |
| 190 | CG9577 RT PCR Rev | CCGATGTCCACCTCCTTGAC |
| 191 | CG17544 RT PCR For | GTGCCCAAGGAGATCGAGAG |
| 192 | CG17544 RT PCR Rev | GTGTTGCTGCCATGCGATAG |
| 193 | CG10467 RT-PCR For | GCCGTATCACACCCGTAGAG |
| 194 | CG10467 RT-PCR Rev | GTTCACGGAGTCGGGAAACT |
| 003 | Rp-49 QPCR For | CCCAAGGGTATCGACAACAGA |
| 004 | Rp-49 QPCR Rev | CGATGTTGGGCATCAGATACTG |
| 115 | EcoR1 3X Flag FASN1 For | CGGAATTCCCATGGACTACAAAGACCATGACGGTGATTATAAAGATCATGACATCGATTACAAGGATGACGATGACAAGCCCGCCCGATTCGCCGAGGA |
| 114 | Xba1 FASN1 cDNA Rev | gatctctagaTTAGTTGAACAGACGCTTCAG |
| 211 | Xba 1 Fatp cDNA 3X Flag Rev | GATCtctagaTTACTTGTCGTCATCGTCTTTGTAGTCGATGTCATGATCTTTATAATCACCGTCATGGTCTTTGTAGTCgaagcggatttcgttgcgctg |
| 209 | Xho1 Fatp 1.8kb cDNA pUASTattB For | GATCctcgagCACCATGGCTGGATTTTTGCTGTGCTCG |

| | | |
|---|---|---|
| 1072 (NS) | hsa miR-125b1 For | ctagcAACATTGTTGCGCTCCTCTCAGTCCCTGAGACCCTAACTTGTGATGTTTACCGTTTAAATCCACGGGTTAGGCTCTTGGGAGCTGCGAGTCGTGCTTTTGCATCCTGGAAg |
| 1073 (NS) | hsa miR-125b1 Rev | CTTCCAGGATGCAAAAGCACGACTCGCAGCTCCCAAGAGCCTAACCCGTGGATTTAAACGGTAAACATCACAAGTTAGGGTCTCAGGGACTGAGAGGAGCGCAACAATGTTGCTAG |
